# Multi-Omics analysis provides crucial insights into ecological adaptation to dryland of a dominant grass (*Psammochloa villosa*, Poaceae) in Northwest China

**DOI:** 10.64898/2026.09.14.751419

**Authors:** Tao Liu, Yuping Liu, Xu Su, Jinyuan Chen, Qian Yang, Rongju Qu, Zhaxi Cairang, Jiarui Jin, Mingjun Yu, Penghui Zhang, Marcial Escudero

**Affiliations:** School of Life Sciences, Qinghai Normal University, Xining, 810008, China; Key Laboratory of Biodiversity Formation Mechanism and Comprehensive Utilization of the Qinghai-Tibet Plateau in Qinghai Province, Qinghai Normal University, Xining, 810008, China; Academy of Plateau Science and Sustainability, Qinghai Normal University, Xining, 810016, China; Plant Biology and Ecology, Faculty of Biology, University of Seville, 41012 Seville, Spain; School of Ecology and Environmental Science, Qinghai Institute of Technology, Xining, 810008, China

**Keywords:** Whole genome duplication, chromosome fusion, gene family expansion, selection sweep, environmental adaptability

## Abstract

Desertification exerts dramatic selection pressures on the evolution of plants. Despite the key role of ecological adaptation by natural selection to arid grasslands and subsequent intraspecific divergence, specific mechanisms driving this process remain poorly understood. *Psammochloa villosa*, a perennial forage grass endemic to the arid grasslands in Northwest China, where it thrives in shifting and semi-fixed sand land due to its exceptional drought tolerance, provides an ideal system to study adaptive evolution to aridity. In our study, we assembled a high-quality, chromosome-scale genome and conducted genomic resequencing of 42 populations across its major distribution. The genome assembly, which is approximately 1.55 Gb in size, has a super-scaffold N50 of 66.79 Mb, with 75.84% of the sequences identified as transposable elements. Coalescent phylogeny and genomic collinearity analyses strongly supported that *P. villosa* and *Neotrinia splendens*, as the closest taxa, shared a recent whole genome duplication (WGD) event occurring approximately 18–20 Mya and followed by their divergence around ∼11.2 Mya. During rediploidization, we identified a chromosomal fusion that reduced the n = 23 chromosomes in *P. villosa* and *Neotrinia splendens* (2n = 46). Based on ancestral grass karyotype (AGK) reconstruction from synteny analysis, our results suggest that, relative to the AGK after the ρ-WGD event, *P*. *villosa* and *N*. *splendens* underwent similar chromosomal restructuring and lineage-specific retention of numerous duplicated copies, providing a genomic basis for potential ecological adaptation and intraspecific diversification. The expanded XTH family, which encodes enzymes mediating xyloglucan endotransglucosylation and hydrolysis and thereby regulating xyloglucan remodeling, showed strong transcriptional responses under PEG-6000 treatment, suggesting that retained copies may be associated with xerophytic adaptation in *P*. *villosa*. Population genomic analyses revealed lineage-specific demographic histories and climate-associated genomic differentiation in *P*. *villosa*. We identified candidate regions enriched for selectively retained duplicated genes. Together, these findings suggest that WGD-derived gene retention created a delayed reservoir of genetic diversity that was later shaped by desertification and Qinghai–Xizang Plateau environmental changes, contributing to climate-associated genomic islands and intraspecific differentiation.

## Introduction

Drought is a critical environmental factor that greatly impacts plant growth and development (Tester & Langridge, 2010). Water demand for agriculture could be double by 2050, while the availability of fresh water is predicted to drop by half, owing to climate change (Gupta et al., 2020). Consequently, drought is anticipated to become more common and severe (Dai, 2012). There is an urgent need to explore drought-resistant plants that use water more efficiently than their present-day counterparts. One potential approach to address this challenge is to decipher the genomic make-up of psammophytes (sand-dune plants adapted to sandy, often unstable substrates characterized by sand burial and shifting substrates), which may be critical for the ecological management and germplasm development of these special wild resources (Wu et al., 2012; Zhang et al., 2022) inhabiting sandy habitats such as fixed or semi-fixed dunes, shifting sand lands, and deserts. However, adaptive genetic resources identified in arid desert-shrub plant in Northwest China are mainly concentrated in a few species: *Populus euphratica* (Zhang et al., 2020), *Nitraria tangutorum* (Wang et al., 2016a), *Capparis spinosa* (Wang et al., 2016b), *Pugionium cornutum* and *Pugionium dolabratum* (Hu et al., 2021). The Inner Mongolia Plateau and adjacent regions harbor heterogeneous and desertified habitats. Most regional plant species have undergone adaptive evolution driven by nature selection such as drought, heat, and sand burial. But studies on the populations’ adaptive evolution of grasses endemic to the area are relatively scarce, which limits the overall understanding of the adaptive evolution mechanism of species in this region.

All naturally occurring grass germplasm potentially harbors a wealth of gene resources (Knapp & Rice, 1994). Dehydration triggers several physiologic and biochemical changes that let grass perceive water-deficiency signals, initiate coping strategies, regulate gene expression, and synthesize key proteins (Koncagül et al., 2018). In water-stressed environments, such essential species must trace suitable locations or adjust to changing environment via phenotypic plasticity or genetic variation. These species are likely to have evolved more adaptable alleles compared to other genetic resources (Aitken et al., 2008; Pardo et al., 2019). Molecular evolutionary investigations of homologs of stress resistance have revealed tandem duplications throughout Pooideae history, drastically boosting their copy number variation and perhaps facilitating drought adaptation (Zhang et al., 2022). In response to the Qinghai-Xizang Plateau environmental changes from the late Pliocene to the Pleistocene, the genomes of two distinct species, *Crucihimalaya himalaica* (Zhang et al., 2019) and *Dactylis glomerate* (Huang et al., 2020), experienced an explosion of long terminal repeat-retrotransposons (LTR-RTs). The absence of key reference genomes has complicated genetic research on grasses due to their high heterozygosity, large proportion of repetitive sequences, and complex co-expression (Copetti et al., 2021; Ren et al., 2022; Zhu et al., 2023).

*Psammochloa villosa* (Poaceae) (2n = 2x = 46; Wang et al., 2021) is a perennial herbaceous plant that is a dominant species endemic of the arid regions of the Inner Mongolian Plateau and the Qinghai-Tibet Plateau in China, and exhibits strong drought tolerance and wind resistance (Lv et al., 2018, 2021). Through long-term adaption to the drought environment, *P. villosa* has developed a considerable level of phenotypic and physiological variation (Liu et al., 2022a). Its root system is critical for anchoring sand in a region where desertification is becoming increasingly problematic (Lu & Kuo, 1987; Dong et al., 1999). As expected, there is abundant genetic diversity in *P. villosa*. Furthermore, altitude and spatial distance are positively correlated with the genetic differentiation coefficient (*F*_ST_), findings consistent with the distribution dynamics based on the spatial distribution modeling (Lv et al., 2021). This herbaceous plant also provides a major forage resource with good palatability and high nutritional value for local livestock in the desert region (Zhang et al., 2003). For a long time, its stalks have also been used as exceptional raw materials for paper making (Yu et al., 2004). We do not yet have a full understanding of this important evolutionary and ecological process. Here, we assembled a chromosome-scale genome for *P. villosa* through various sequencing strategies involving Illumina sequencing, PacBio HiFi sequencing reads and chromosome conformation capture (Hi-C) technology. We conducted comparisons among Poaceae genomes and estimated phylogenies and divergence times between *P*. *villosa* and *N. splendens* around 10-12 Mya. Late Miocene stepwise aridification in the Asian interior around 18-20 Mya, driven by global cooling and Tibetan Plateau uplift, created expanding deserts and grasslands. And, demographic history analysis also revealed that *P*. *villosa* has undergone a first bottleneck during the aridification stage (7–10 Mya, Wen et al., 2023). We detected genetic differentiation among key lineages by quantified polymorphism levels (*θπ*), genetic differentiation (*F*_ST_), and selection statistics (Tajima’s *D*) and identified the selected region. Based on gene-environment associations (GEA) in genome-wide studies, collinear gene analyses, structural variation of chromosomes and RNA-seq-based transcriptomics, we aimed to clarify the genomic basis of how this species ecological adapts to drought habitats.

## Materials and methods

### 2.1 Plant materials and growth conditions

We collected seeds of *P. villosa* from 2016 to 2021 in the northwestern regions of China from 153 different populations spread over five provinces (Table S1). To make a more accurate comparison among the biological replicates, we attempted to grow 50 fully developed seeds from a single population (P19-037 of Table S1). Following the procedures outlined by Kim et al. (2009), we stored the mature seeds in the laboratory at 4°C in the laboratory until planting them. We surface-sterilized the seeds before sowing by immersing them in 15% (v/v) sodium hypochlorite solution for 15 min (Dkhil & Denden, 2010). To ensure consistent germination conditions, we allowed seed germination in culture dishes for a week under dark conditions and moistened with distilled water. After that, we grew seedlings in a thermostatic incubator (28/22°C) with a 16-hour light/eight-hour dark cycles and 60% relative humidity. Following Liu et al. (2022a), we planted three seedlings in 0.5 kg soil pots and irrigated them with 100 mL Hoagland nutritional solution every three days.

### 2.2 DNA extraction, Genome, Transcriptome and Hi-C sequencing

For genome sequencing, we chose one of the best-growing individuals (P19-037, Table S1) based on its growth state under normal seedling cultivation light. We harvested its leaves, immediately froze them in liquid nitrogen, applying a modified CTAB method to extract genomic DNA from healthy leaves, stems and roots (Fang et al., 2021). RNase was used to remove RNA contaminants. Following DNA extraction, we assessed the quality by agarose gel electrophoresis. Genome size was estimated by combining flow cytometry and whole-genome survey sequencing. Using flow cytometry (Partec CyFlow Space, Germany) on propidium iodide–stained nuclei isolated from young leaves (following a modified Doležel and Bartoš protocol), we previously estimated the genome size as 1580.10 ± 5.02 Mb (Liu et al., 2022).

For the whole-genome survey sequencing, a PE150 short-read library was constructed and sequenced on the Illumina NovaSeq 6000 platform from high-quality DNA. *K*-mer analysis was performed using Jellyfish (Marcais & Kingsford, 2011) generated 42,100,563,004 distinct 17-mer sequences. The genome size was calculated using the formula (Genome size=*K*-mer_number/Peak_depth) based on a 17-mer depth of 13, and heterozygosity was estimated from the *K*-mer Poisson distribution pattern (Figure S1). Using this sequencing-based approach, we estimated the genome size of *P*. *villosa* to be 1564.195 Mb (Table S2) with GenomeScope v2.0.1 (Ranallo-Benavidez et al., 2020).

To obtain high-resolution haplotype assembly for the *P. villosa* genome, we isolated a total of 30 μg of DNA (with A260/280=1.8-2.0 and A260/230⩾2) from 15 g fresh leaf tissue (P37, Table S1) and performed PacBio HiFi sequencing on the Sequel II instrument. HiFi sequencing across three SMRT cells generated over 20× genomic coverage, yielding long reads (median of CCS length distribution over 15 kb) and highly accurate (Q20 > 99%) HiFi reads (Guan et al., 2020). Using about 2g the freshest leaves from the same individual of the HiFi library and for material of a Hi-C library, we performed the Hi-C pipeline described in previous studies (Burton et al., 2013; Rao et al., 2014). The detailed procedures for its implementation are provided as follows. (1) We cut the leaves into 1- to 2-mm strips, which were fixed with fresh formaldehyde at a final concentration of 2% in NIB buffer (20 mMol/L HEPES Solution, pH 8.0, 250 mMol/L sucrose, 1 mMol/L MgCl_2_, 5 mMol/L KCl, 40% (v/v) glycerol, 0.25% (v/v) Triton X-100, 0.1 % (v/v) PMSF Solution, and 0.1% (v/v) β-mercaptoethanol) at 4℃ for 45 min in a vacuum (Rao et al., 2014). Under vacuum infiltration, we used 0.375 Mol/L final concentration glycine to quench the formaldehyde for an additional 5 min. We washed the sample twice by ice-cold water. The clean samples were ground to powder in liquid nitrogen and resuspended in NIB buffer and filtered with one layer of Miracloth. (2) The isolated nuclei were lysed with 0.1% (w/v) final concentration SDS at 65℃ for 10 min and then SDS molecules were quenched by 1% (v/v) final concentration Triton X-100. (3) The DNA in the nuclei was digested by the four-cutter restriction enzyme, MboII at 37°C for 2 hr (Rao et al., 2014). (4) We labeled restriction fragment ends with biotinylated cytosine nucleotides using biotin-14-dCTP (TriLINK) to blunt the sticky ends resulting from digestion, biotinylate these ends, and facilitate their subsequent capture via biotin beads. (5) Blunt-end ligation was carried out at 16°C overnight in presence of 50 Weiss units of T4 DNA ligase (Crane et al., 2015). (6) After ligation, the cross-linking was reversed by 200 µg/mL proteinase K (Thermo) at 65°C overnight. Furthermore, DNA purification was achieved through QIAamp DNA Mini Kit (Qiagen) according to manufacturers’ instructions. And purified DNA was sheared to a length of 400 bp. Point ligation junctions were pulled down by Dynabeads® MyOne™ Streptavidin C1(Thermofisher) according to manufacturers’ instructions. Finally, the Hi-C library for Illumina sequencing was prepped by NEBNext® Ultra™ II DNA library Prep Kit for Illumina(NEB) according to manufacturers’ instructions. Fragments between 400 and 600 bp were paired-end sequenced on the Illumina HiSeq X10 platform platform (San Diego, CA, United States) with 150 PE mode. Three replicates were generated for one group material.

To achieve non-redundant and high-confidence genomic annotation, we established transcriptome libraries, including a full-length transcriptome library of the natural population and nine short-read libraries from seedlings. Samples of fresh tissue (leaves, stems, and roots) of *P. villosa* were taken from the species’ natural habitat in China’s Ordos City of inner mongolia (same population as seedlings grown for genomic sequencing; Table S1). To increase the accuracy of genome annotation, we kept mixed tissue to extract total RNA, and stored the collection in liquid nitrogen in the field and at -80°C in the lab until processing. We extracted RNA using a Dynabeads mRNA DIRECT Kit from Thermo Fisher Scientific, Inc. We measured RNA concentration and quality using Agilent Bioanalyzer 2100 and Qubit (Thermo Fisher Scientific, Inc.). We constructed two ISO-Seq libraries by using about 2 *μ*g of total RNA for cDNA synthesis and an optimized SMRTbell Template Prep Kit 1.0. We used the SMRT Cell method from Pacific Biosystems to sequence with two cells (GENEWIZ Co., Ltd, Suzhou, China).

For the transcriptome profile analysis of drought-related gene families, we germinated seeds from the same population as those grown for the PacBio full-length cDNA sequencing. These seedlings with the same growth were treated with 0%, 10%, and 20% PEG-6000. We extracted total RNA from 0.5 mg of the mixed tissues stored in liquid nitrogen using a DP432 RNA extraction kit (TIANGEN Biotech, Beijing). After quantifying the extracted RNAs, RNAs with polyA tail were purified using beads with oligonucleotides (dT) and fragmented into short sequences that we used as templates for cDNA synthesis (Mommaerts et al., 2015). We constructed nine different cDNA libraries, three biological replicates for each group, each with a 350 bp insert size, and sequenced them on a NovaSeq 6000 (Beijing Biomarker Technologies Co.), with each library generating over 20 million read pairs (Table S3).

### 2.3 Genome assembly for chromosomal construction

We applied the filtered HiFi reads to perform *de novo* assembly contigs using hifiasm v0.16.1-r375 (Cheng et al., 2021), and identified and removed haplotypic duplications with purge_dups software v1.2.5 (Guan et al., 2020) to obtain accurate contigs sequences. Subsequently, we enhanced the draft contigs by mapping all filtered PacBio reads to the assembly using pbmm2 (with --preset SUBREAD for subreads or --preset CCS for HiFi/CCS reads and --sort enabled), and then generated an Arrow consensus using gcpp (--algorithm=arrow) from the PacBio SMRTLink suite (Chin et al., 2013) to improve assembly quality of the genome.

A total of 339 Gb of raw data consisting of Hi-C read pairs was initially aligned to the scaffolds using BWA mem (version 0.7.17, Li & Durbin, 2009) with the default parameters. Only the valid interaction pairs were used for constructing the interaction map of genomes. These retained reads were clustered and anchored to the scaffold onto 23 pseudochromosomes by Juicer v2.0 (Durand et al., 2016b). We further filtered the assembly outcomes and manually corrected construction errors visually using the Juicebox tool v1.8.8 (Durand et al., 2016a). These errors included misjoins, translocations, inversions, and chromosomal boundary errors (Burton et al., 2013). After that, we used 3D-DNA v.1.8.09 (Dudchenko et al., 2017) to construct the interaction matrix for JuiceBox Assembly Tools (JBAT), representing the assembly with contigs organized into candidate chromosomes. Using JBAT, we interactively updated contigs’ location, orientation, and chromosome delineation. Finally, the genome assembly of *P. villosa* was organized into 23 chromosomes supported by Hi-C reads.

We evaluated the quality of the final genome assembly in terms of contiguity, completeness, and sequence concordance. Contiguity was assessed using the N50 value and QUAST software (Quality Assessment Tool for Genome Assemblies, Version 5.2.0, Gurevich et al., 2013). completeness was evaluated using BUSCO (Benchmarking Universal Single-Copy Orthologs, v5.12, Simão et al., 2015) with the embryophyta odb9 dataset (1,440 conserved single-copy orthologs) and LAI value (the LTR Assembly Index, Ou et al., 2018); and sequence concordance was examined by mapping full-length transcripts from our previous study (PRJNA730046) to the final assembly using GMAP (Genomic Mapping and Alignment Program, gmap.avx2 v2021-12-17) with default parameters (Wu & Watanabe, 2005), where a mapping rate >90% and coverage >95% indicated good agreement between the assembly and transcriptomic data.

### 2.4 Annotation of repetitive elements, non-coding RNAs (ncRNAs)

In the genome of *P. villosa,* we made predictions regarding repetitive elements such as tandem repeats and transposable elements (TEs). We used TRF (Tandem Repeats Finder, version 4.09, Benson, 1999) to look for tandem repeat sequences, and the parameters for the program were as follows: Match = 2, Mismatch = 7, Delta = 7, PM = 80, PI = 10, Minscore = 50, MaxPeriod = 2000. The annotation of interspersed repeats, such as TEs, was accomplished primarily by integrating homology-based and *ab initio* predictions. For homolog annotation, we adopted RepeatMasker v.4.0.5 (Smit, 1996) with the Repbase database (Poales) and the RepeatProteinMasker v4.1.4 (Tarailo-Graovac & Chen, 2009) as the local TE library (Jurka et al., 2005). We made consolidation and filtration from LTR harvest (module of GenomeTools software version 1.6.2, Ellinghaus et al., 2008), LTR_FINDER v1.2 (Xu & Wang, 2007), and LTR_retriever v2.9.0 (Ou & Jiang, 2018) to search Long terminal repeat-retrotransposons (LTR) and analyze insert time of the genome. MITE-Hunter (running Perl version 5.16.3, Han & Wessler, 2010) was used to finder DNA MITE of *P. villosa*. After that, we used *de novo*-based identification with RepeatModeler version 2.0.3 (http://www.repeatmasker.org) to predict the repeat elements and family relationships of the genomic data. We processed alignment to the transposase protein database for unclassified repeated sequences and utilized DeepTE v1.0.6 (set -m Plants_model from DeepTE model and -fam LTR, Yan et al., 2020) for further identification. We produced the final annotation of repeats by integrating all of the results of repeat identification produced by the various software and eliminating redundancy through a custom script (https://github.com/Liutao93/Pvi_genome.git).

We predicted transfer RNAs (tRNAs) using tRNAscan-SE v1.3.1 (Lowe & Eddy, 1997) with eukaryotic parameters. We annotated rRNA, including 5.8s, 18s, and 28s, in the genome via RNAmmer v1.2 software (Lagesen et al., 2007). Additionally, we used Infernal cmscan v1.1.1 (Nawrocki & Eddy, 2013) to search the Rfam database v12.0 (Nawrocki et al., 2015) for non-coding RNA (ncRNA) sequences, and identified various types of ncRNAs, such as microRNA and small nuclear RNA (Hunter et al., 2008; Nawrocki & Eddy, 2013).

We predicted protein-coding genes using a combination of *de novo* prediction, homology-based, and transcriptome-based prediction. After masking the repeat sequences in the genome, we trained the gene models of *P. villosa* using a Hidden Markov Model and randomly selected 3000 full-length genes from the transcriptome-based predictions made by PASA v.2.5.2 (Haas et al., 2003). We processed the alignment of all full-length transcripts to the genomic sequence and generated transcripts assembly from methods of Hisat2 v2.2.1 and StringTie v2.2.1, respectively. For *de novo* prediction, the gene model of *P. villosa* was trained by MAKER2 (version 2.31.11, Holt & Yandell, 2011), which incorporates the Augustus version 3.3.2 program (Stanke et al., 2006), the GeneMarke-ET version 3.54 program (Stanke et al., 2008), and further predicted by SNAP version 2013_11_29 (Korf, 2004), the GlimmerHMM version 3.0.4 program (Majoros et al., 2004), and the GeneID version 1.4.5 program (Alioto et al., 2018).

In addition, we built the RNA library from mixed samples (root, stem, and leaf) to make predictions based on the transcriptome. Trinity software version 2.1.1 (Grabherr et al., 2011) was used to *de novo* assemble of the RNA-seq reads into transcripts. Together with the complete full-length transcript, we produced the annotation file using the PASA pipeline version 2.5.2 to predict the protein-coding genes (PCGs), and then TransDecoder v.5.5.0 (http://transcoder.github.io) to eliminate incorrect gene structure. For homology-based predictions, GeMoMa version 1.3.1 (Keilwagen et al., 2016) was utilized, and it was fed protein sequences from a total of eight distinct species of organisms (*Neotrinia splendens, Arabidopsis thaliana, Brachypodium distachyon, Cleistogenes songorica, Oryza sativa, Sorghum bicolor, Setaria italica* and *Zea mays*). Finally, based on the EVidenceModeler (EVM) software version 1.1.1, all of the gene predictions produced by the methodologies mentioned above were incorporated into a consensus set of genes (Haas et al., 2008).

Functional annotations of predicted protein-coding genes were obtained using two complementary approaches: sequence similarity–based annotation, performed with eggNOG-Mapper version 2.1.12 (http://eggnog-mapper.embl.de/) and BLASTP searches (BLAST+ version 2.12.0; Camacho et al., 2009) against publicly available protein databases, (UniprotKB, GO, KEGG, and NR; Boeckmann et al., 2003; Kanehisa & Goto, 2000) with an e-value of 1e-5; and conserved domain–based annotation, performed using InterProScan version 4.8.6.8 to search the InterPro, SMART, and Pfam databases (Zdobnov & Apweiler, 2001).

### 2.5 Gene family clustering and phylogenetic analysis

To investigate the evolution and divergence of the genome of *P. villosa*, we collected protein-coding gene sequences from eleven different species of Poaceae (*Neotrinia splendens*, *Aegilops tauschii*, *Brachypodium distachyon*, *Hordeum vulgare*, *Oryza sativa*, *Oropetium thomaeum*, *Pharus latifolius*, *Phyllostachys edulis*, *Sorghum bicolor*, *Setaria italica* and *Zea mays*), one specie of monocotyledon from Orchidaceae (*Phalaenopsis equestris*), and one eudicot from Brassicaceae (*Arabidopsis thaliana*) for comparative analyses. We extracted the longest transcript of each gene from each genome, disregarding alternative splicing (Yin et al., 2020), and calculated pairwise sequence similarities based on all-versus-all BLASTP using the default settings (e-value 1e-5 and a minimum match length of 50%). We used OrthoMCL version 2.0.9 to cluster the BLASTP results and divide them into orthologs and paralogs (Li et al., 2003). We used a Venn diagram to illustrate the percentages of gene families shared between closely related Poaceae species and those unique to each species (*Neotrinia splendens*, *Orinus kokonorica*, *Oryza sativa*, *Psammochloa villosa* and *Zea mays*) via a Perl script (https://github.com/Liutao93/Pvi_genome.git). We identified single-copy orthologous genes from the clustering results and performed multiple alignments using MUSCLE version 3.8.31 (Edgar, 2004) with the default parameters. We removed regions with poor alignment or large differences using Gblock version 0.91 (Talavera & Castresana, 2007). Then, using the alignment result, we concatenated those sites model to produce a super-gene matrix. We reconstructed the phylogenetic tree using RAxML v8.1.17 (Stamatakis, 2014) with the best JTT + I + G + F model according to AIC and BIC values from ProtTest v.3.4.2 (Darriba et al., 2011), performed 200 bootstrap replicates. Finally, we visualized the resulting phylogeny in iTOL v.3.0 (https://itol2.embl.de/itol.cgi; Letunic & Bork, 2021).

We estimated the species divergence times by calculating the average substitution rate of *P. villosa* along each branch using the baseML module and the MCMCtree module in PAML v.4.5 (Yang et al., 2007), with the "correlated rates clock (clock = 3)" model and the "F84" substitution model. To further calibrate the divergence times, we applied the divergence times for *A. thaliana*-*O. sativa* (mean: 160.0 Mya, std dev: 4.0) as calibrations for the divergence times obtained from the TimeTree tools (http://www.timetree.org; Clarkson et al., 2017). To precisely correct the divergence time between *O. sativa*-*P*. *villosa* (Pooideae and Oryzoideae), we used two fossil calibrations of rice tribe (Oryzeae). Two newly phytoliths fossils, *Changii indicum* and *Tateokai deccana*, were placed at the base of Oryzeae within Ehrhartoideae and were used to calibrate the divergence between Oryzoideae and the other lineages at 72.38 Mya (calibrated herein 95% HPD: 61.40–83.67 Mya; Prasad et al., 2011).

### 2.6 Estimation of Genome synteny and whole genome duplication

We performed homologous alignments both within and among the genomes of, *O. sativa*, and *P. villosa*. We detect putative homologous genes for accurate genomic collinearity analysis and, consequently, to filter out redundant or mismatch information. In addition, we adopted MCscanX (Wang et al., 2012) and the CollinearScan (Wang et al., 2006) to infer homologous blocks involving collinear genes both within and between genomes. We constructed homology dotplots for inter-species comparative genomes using a custom Python script (https://github.com/Liutao93/Pvi_genome.git) and set a maximum gap length of 50 genes between collinear genes along a chromosome region, following Wang et al. (2017, 2018). We determined the nonsynonymous (*K*a) and synonymous (*K*s) substitution rates for gene pairs using *K*a*K*s_Calculator v2.0 (Wang et al., 2010) under a YN model and converted protein alignments into codon alignments via PAL2NAL v14 (Suyama et al., 2006). Based on the synteny analysis of cross-species genomes, we extracted complete synteny blocks (Tang et al., 2008) and used the Whole-Genome Duplication Integrated analysis tools (WGDI v0.75, Sun et al., 2022) to facilitate the reconstruction of detailed karyotype evolution.

### 2.7 Identification of gene families’ expansion and contraction and related with drought tolerance

We regarded of the orthologous clustering results from OrthoMCL as input for the Computational Analysis of gene Family Evolution (CAFE) version 5.0 (De Bie et al., 2006) to obtain gene families along each branch of the phylogenetic tree using birth-and-death mode. Using conditional likelihood as the test statistic, we estimated the corresponding *P*-values of each lineage and inferred significant expansion or contraction of gene families with *P*-values of 0.01 as the threshold. Functional enrichment analysis of gene ontology (GO; Ashburner et al., 2000) was carried out to discover the functions of both the expanded and the contracted gene families.

To identify genes involved in drought tolerance in *P. villosa*, we used both BLAST alignment against the *Arabidopsis* Information Resource (TAIR) database and a hidden Markov model (HMM) from the Pfam database. We searched all protein sequences annotated in the genome using HMMER v.3.2.1 for Pfam database (Finn et al., 2011; 2016) analysis to locate homologous gene families. Then, we manually identified genes and transcription factors (TF) involved in regulating drought tolerance in *P. villosa* using PlantTFDB version 4.0 (Jin et al., 2017). Only genes with identical protein domains were chosen as orthologs.

To investigate the genome-wide responses to drought tolerance in *P. villosa*, we conducted transcriptomic analysis on mixed root, stem, and leaf tissues treated with 10% and 20% PEG-6000 for 48 hours. To obtain non-redundant clean data, we removed adapters and low-quality reads from the raw sequencing reads using FASTP version 0.20.0 (Chen et al., 2018) with phred quality ⩾ Q10 and 50% of bases unqualified, if one read’s number of N base over 10_base_limit, then discarded this read pair. We mapped the transcriptome data onto the genome using HISAT2 (Daehwan et al., 2015) and assembled transcripts with StringTie v1.3.1 (Pertea et al., 2015). The counts of mapped reads determined the TPM value of each gene. We analyzed differential expression using DESeq2 (Love et al., 2014), identifying differentially expressed genes (DEGs) as those with expression levels altered by at least twofold compared to the control group (false discovery rate, FDR ≤ 0.05). We performed a *K*-means clustering analysis of DEGs using the OmicShare tools and conducted a KEGG enrichment analysis on the genes within each cluster.

### 2.8 Genome resequencing and variant calling and filtering

To reveal population genomic variation of *P. villosa* and climatic association, we surveyed 153 wild populations across the species’ entire distribution area in the Inner Mongolian Plateau and its adjacent regions from 2016 to 2021, from which we selected 42 representative populations (three individuals per population; 126 individuals in total) for whole-genome resequencing.

These 126 individuals were resequenced on the BGI T7 platform using 2×150 bp paired-end reads at > 10× resequencing depth. Raw data of resequencing were quality-filtered using FASTP (v0.20.0). We removed adapter sequences of the raw data by providing the adapter FASTA file supplied by the sequencing facility and specifying the --adapter-fasta option. Also, we filtered low-quality reads by using –qualified-quality-phred as 10, --unqualified-percent-limit as 50, and –n-base-limit as 10. The resulting clean paired-end reads were aligned to the *P*. *villosa* reference genome using BWA-MEM v0.7.17 (Li & Durbin, 2009) with default parameters. The resulting alignments were converted and merged using SAMtools v1.1 (Li et al., 2009), sorted with SortSam of Picard, and removed duplication with MarkDuplicates of Picard (v1.92; http://broadinstitute.github.io/picard/). We then performed variant calling using the Genome Analysis Toolkit (GATK v3.8; McKenna et al., 2010). The InDel intervals were realigned using GATK to enhance alignments near InDels. Variants were called in each accession separately using haplotypecaller module. Individual genome variant call format (GVCF) files were merged using genotype gvcfs module. After that, the variants were classified as single nucleotide polymorphisms (SNPs) and InDels using Shell script (https://github.com/Liutao93/Pvi_ Pvi_ReSeq.git). These variants were initially filtered using VariantFiltration (for SNPs: --filterExpression "QD < 2.0, FS > 60.0, MQ < 40.0, MQRankSum < −12.5, ReadPosRankSum < −8.0"; for InDels: --filterExpression "QD < 2.0, FS > 200.0, ReadPosRankSum < −20.0") of GATK. Furthermore, the SNPs at or within 5 bp from InDels were also removed using VCFutils (vcfutils.pl; distributed with SAMtools v1.1). The SNPs were further filtered using vcftools (v0.1.13) with the following parameters: maf < 0.05, min--meanDP =3, max-meanDP = 27, max-missing = 0.7, hwe = 0.001, minGQ = 10. A final data set with a total number of 8,774,438 high-quality SNPs was generated across 126 individuals and was subsequently functionally annotated against the *P*. *villosa* reference genome using ANNOVAR (Perl 4.3; table_annovar.pl) for downstream analyses.

### 2.9 Population genetic structure, selective sweep and gene flow

We calculated the genetic distance matrix using PLINK (v1.90b3y; Purcell et al., 2007) and then performed principal component analysis (PCA), maximum-likelihood (ML) phylogenetic analysis based on the distance matrix, and population structure inference using ADMIXTURE (v1.23; Alexander et al., 2009) by testing *K* = 2–10; the optimal *K* was determined as the one with the lowest cross-validation (CV) error, and the inferred ancestry proportions and admixture patterns were visualized geographically using the R package mapmixture (Jenkins, 2024) in combination with ArcGIS version 10.3 (Esri, 2014). We used TreeMix (v0.5.4; Pickrell and Pritchard, 2012) to infer population divergence history and gene flow by fitting a maximum-likelihood tree to allele-frequency data, rooting *N*. *splendens* (From the COMMSBIO-25-4409 variant dataset) as an outgroup. We treated the observed allele-frequency covariance among populations as the real value and the expected covariance under the relationship within ML tree as the estimated value, and inferred gene flow when the real value was substantially lower than the estimated value (i.e., the tree would otherwise overstate divergence), after which TreeMix then added migration edges to explain these residual patterns and infer the direction and magnitude of admixture. We ran TreeMix with the -global, -se, and -noss options to model genome-wide drift, estimate standard errors of migration weights, and avoid overcorrection for small sample sizes.

### 2.10 Genetic diversity, differentiation and demographic history

We inferred genetic diversity, selective sweep, linkage disequilibrium for further analysis by the level of nucleotide diversity (*θπ*) varied across the different populations, Genetic divergence (*F*_ST_), and the reduction of diversity (ROD) with a 50 kb nonoverlapping window (Danecek et al., 2011). We combined an empirical approach with a permutation approach to identify outlier windows, which genes under positive selection. We performed the cross-population composite likelihood ratio test (XP-CLR) and Tajima’s *D* value to detect decreased levels of genetic diversity under selective sweeps across the genome (Chen et al., 2010), measured and compared patterns of linkage disequilibrium (LD) decay for different groups using the software LDdecay version 3.40 (Zhang et al., 2019). Demographic history and effective population size fluctuation over time for the eastern and western lineages inferred by population structure analyses were inferred using the pairwise sequential Markovian coalescence (PSMC) model (v0.6.4-r49) (Li & Durbin, 2011). The analysis was performed using the following parameters were −N25 − t15 − r5 – p “4 + 25 × 2 + 4 + 6”. A generation time of five years (Vintsek et al., 2022; Jin et al., 2025), given that the species is perennial and a substitution rate (*μ*) of 1.3e-8 per site per year, which was estimated by PAML program. SMC++ (Terhorst et al., 2017) was used to infer the demographic history in the recent past and the split time of the two lineages from the common ancestor. We randomly selected 18 individuals from the eastern middle and western groups. SMC++ analysis was performed twice based on two replicated random selections from the three lineages.

We conducted the Hudson-Kreitman-Aguadé (HKA) test comparing polymorphism (*π*) and divergence (*D_xy_*) between candidate regions and neutrally evolving loci to infer signatures of selection. The HKA statistic was calculated using custom Perl scripts based on the Hudson-Kreitman-Aguadé framework (HKA test v2.0; https://github.com/popgen-tools/hka). The statistical significance of HKA test was determined by 10,000 coalescent simulations under a neutral model. Observed ratios exceeding this distribution defined *P*-values, with regions showing *P* < 0.05 deemed to exhibit significantly different mutation patterns from background levels.

Demographic histories of there lineages were inferred using the continuous-time sequential Markovian coalescent approximation implemented in fastsimcoal2 v2.7.0.9 (Excoffier et al., 2021). The observed joint site frequency spectrum (SFS) was constructed from SNPs dataset of 126 individuals using easySFS (github.com/isaacovercast/easySFS), projecting the data to a uniform sample size to account for potential biases. Guided by TreeMix analysis indicating a single major gene flow event, competing demographic models were designed to test scenarios of divergence with and without migration among the three identified groups. Parameter estimation was performed using the expectation-conditional maximization (ECM) algorithm, with each model undergoing 100 independent optimization runs (-n 100,000 -L 40). The best-fit model was selected based on likelihood scores, and parameter uncertainty was assessed using 100 parametric bootstrap replicates. Absolute divergence times and migration rates were estimated using a mutation rate of 1.3e-8 per site per year.

### 2.11 Identification of environment-associated genetic variants

We used two complementary genotype-environment association (GEA) approaches to detect the environment-associated genetic variants. First, we tested for GEAs for 19 abiotic variables (https://worldclim.org/), water vapor pressure (vapr, kPa) of twelve months and wind speed (wind, m·s^-1^) of twelve months using the latent factor mixed model (LFMM v2.0, Fick & Hijmans, 2017), which tests for associations between genotypes and environment variables while accounting for background population structure (R package named LEA, R version 4.2). We also used a complementary multivariate landscape genomic method, redundancy analysis (RDA) to identify co-varying variants that were likely associated with multivariate environment predictors. Significant environment associated variants were defined as those having loadings in the tails of the distribution using a standard deviation cutoff of 3 along one or more RDA axes. Based on the ancestry clusters from ADMIXTURE, we inferred population structure in the genotype data with three latent factors. For each environmental variable, we ran five independent MCMC runs using 5000 iterations as burn-in followed by 10,000 iterations. *P* values from all five runs were then averaged for each variant and adjusted for multiple tests using a false discovery rate (FDR) correction of 5% as the significance cutoff. Second, after considering the ranked importance of the 19 environmental variables estimated using GF analyses with R package “gradient Forest” and correlations among the variables, four environmental variables (BIO10, BIO15, BIO18 and vapr12) with pairwise correlation coefficients |*r*| < 0.6 were selected for the RDA analyses using the R package (Vegan 2.6-4).

## 3 Results

### 3.1 Genome assembly, quality assessment and annotation

*Psammochloa villosa* is a diploid plant with 2n = 2x = 46 chromosomes. To sequence and assemble the genome of *P. villosa*, we constructed the PacBio CCS library (HiFi), a 150 bp paired-end genomic library with 300 bp insert sizes, a Hi-C library, and PacBio SMRT cells for the full-length transcriptome. A total of 69.81 Gb of raw data (over 20× genome coverage) were produced using the PacBio Sequel II technology, and 83.49 Gb of short reads (over 50× genome coverage) from the Illumina HiSeq platform were generated, indicating adequate fold coverage of the *P. villosa* genome (Table S3). To assign scaffolds to the correct chromosomal positions, a total of 129.8 Gb Hi-C reads (∼100x genome coverage) were generated based on genetic proximity. We successfully anchored 98.82% of the complete contigs assembly into 23 pseudochromosomes (Figure S2 and Table S4). In the genome of *P. villosa*, sixteen pseudochromosomes out of the 23 had no gaps. The final assembly length of the genome was 1522.89 Mb, and the N50 length of the super-scaffold was 66.80 Mb, which is somewhat lower than the previously predicted genome size, perhaps due to the genome’s relatively high heterozygosity and repeat proportion.

To correct long-read sequencing errors, we mapped paired-end short reads to the genomic assembly, resulting in 97.95% of short reads being successfully mapped to the genome. Furthermore, the 36.47 Gb of PacBio full-length transcriptome data generated 184,076 full-length isoforms (Table S5), of which around 98.95% (73,770) successfully matched to the genome (Table S6). We assessed the completeness of the assembled genome using BUSCO and the Poales gene set (odb10) and found that 4738 out of 4896 (96.8%) conserved protein genes were entirely captured in the genome (Table S7). The LAI score of *P. villosa* was determined to be 20.98, demonstrating the comprehensive advantages of using genome and transcriptome long-read-based sequencing techniques (Table S8).

### 3.2 Long terminal repeat retrotransposon classification and genes annotation

Using *de novo* and homology-based approaches, we analyzed and annotated repetitive sequences and transposon elements (TE), including long terminal repeat retrotransposons (LTR-RTs), MITE as well as small class non-autonomous TEs. Approximately 1207 Mb, accounting for 75.84% of the assembled genome, was identified as DNA transposons, retroelements, satellites, and simple repeats (Table 1). LTR-RTs were the most abundant repetitive sequences in the genome, accounting for 56.23%, with *Gypsy* and *Copia* being the two prevalent LTR-RT families, accounting for 17.13% and 39.30% of the genome assembly, respectively (Figure 1). The *Gypsy*/*Copia* elements ratio of *P*. *villosa* (2.29) showed a similar LTR retrotransposon composition pattern to other Poaceae genomes (Table S8). We estimated that LTR-RTs may have undergone expansion in the grass genome during the last one millon years (Figure S3).

**Figure 1.**
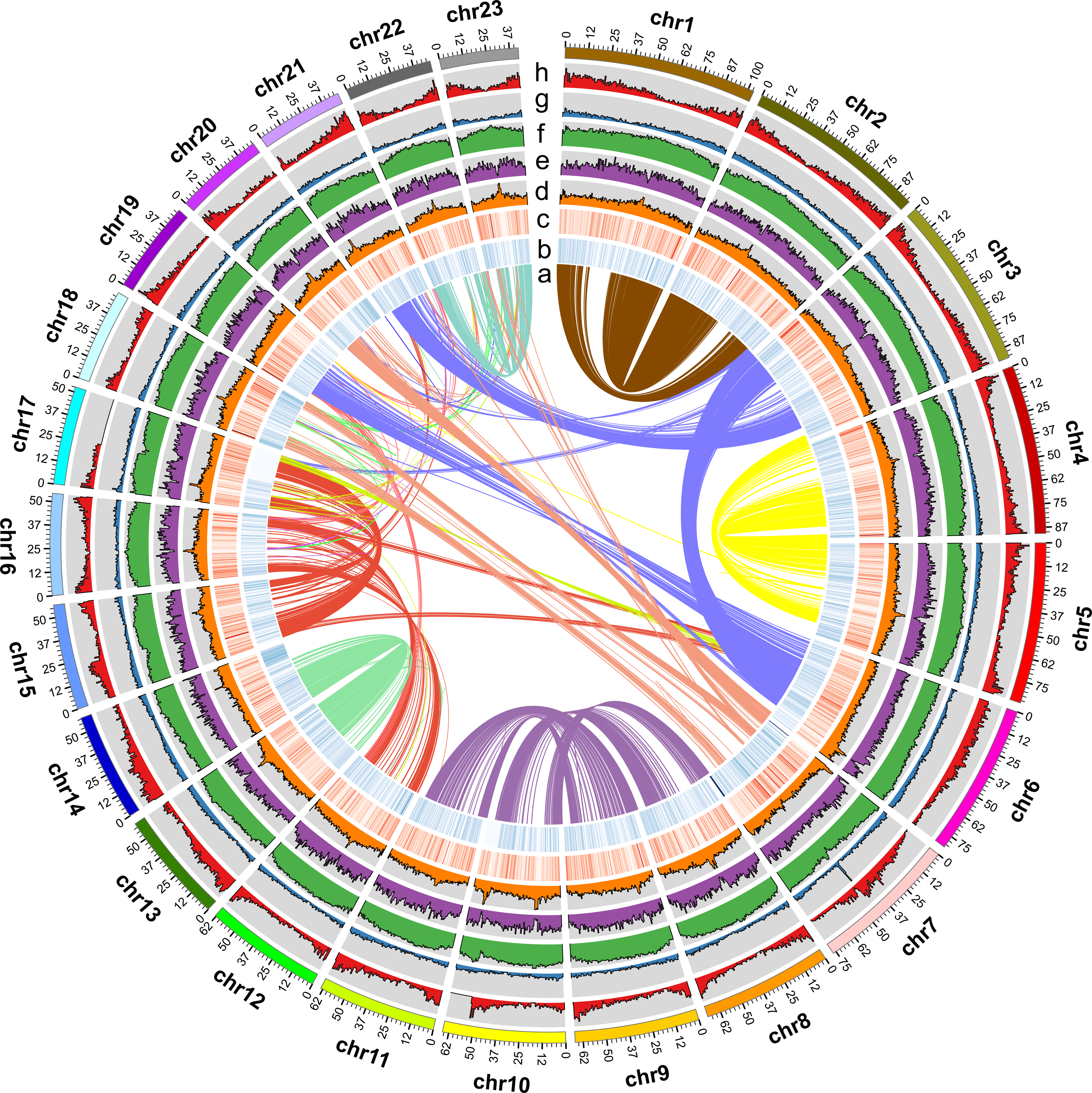
The overview genome assembly of *P. villosa*.

By integrating annotations from *ab initio*, homology-based, and transcriptome-based methods, we predicted 54,317 protein-coding genes with high confidence (Table S9). Most of these genes (98.795%) were successfully anchored onto one of the 23 pseudochromosomes. The length of protein-coding genes in the genome of *P. villosa* was 3,250 base pairs on average and spanned 4.85 coding exons (CDS only; excluding 5′/3′ UTRs), similar to those found in other Poaceae species (Figure S4). We identified 51,492 genes with functional annotations in protein databases, representing 94.79% of the predicted genes (Table S10). Additionally, we discovered 1554 small nuclear RNA (snRNA) genes, 542 microRNA (miRNA) genes, 1089 transfer RNA (tRNA) genes, and 6221 ribosomal RNA (rRNA) genes in the genome (Table S11). We concluded that the genome of *P. villosa* has a high repetition rate based on genomic annotation.

### 3.3 Genome evolution, phylogenetic analysis in Poaceae

To gain insight into the evolutionary history of *P. villosa*, we compared its genome with the publicly available genomes of 14 species. Based on homolog searches, we found that 12 species of Poaceae shared 119,005 gene families (Figure S5), in which 3,203 families were unique to *P. villosa* and *A. tauschii* and *N. splendens* (Table S12). These unique gene families were enriched in "guard cell differentiation", "response to oxidative stress, water", "peroxidase activity" and "xylosyltransferase activity " GO categories (Table S13). We further detected 48 expanded and 42 conserved (Fisher’s exact test, *p* < 0.01) gene families in the genome of *P. villosa* when compared with other 11 Poaceae species. GO enrichment analysis of these gene families revealed significant relevance in "guard cell differentiation ", " stomatal complex development", "plant epidermal cell differentiation" and hydrogen peroxide catabolic process"(Figure S6).

To reconstruct the phylogeny of *P. villosa*, we identified 294 single-copy orthologs among 14 species, using *Arabidopsis thaliana* (Family: Brassicaceae) and *Phalaenopsis equestris* (Family: Orchidaceae) as outgroups. Phylogenetic analysis revealed that *Oryza sativa* was in the same lineage as the other six species that were classified into the BOP clade (*P. edulis*, *N. splendens*, *P. villosa*, *B. distachyon*, *H. vulgare* and *A. tauschii*). Notably, *P. villosa* and *N. splendens* formed a subclade (Tribe: Stipeae) that was phylogenetically sister group to the clade comprising *B. distachyon* (Tribe: Brachypodieae), *A. tauschii* and *H. vulgare* (Tribe: Triticeae). Using MCMCtree and fossil calibration, we determined that the divergence between *P. villosa* and *N. splendens* occurred approximately 7.17 Mya (Figure 2, 95% HPD: 2.277-13.946). The phylogenetic divergence between *B. distachyon* and the subclade formed by *A. tauschii* and *H. vulgare* was estimated around 32.02 Mya (95% HPD: 19.735-44.656).

**Figure 2.**
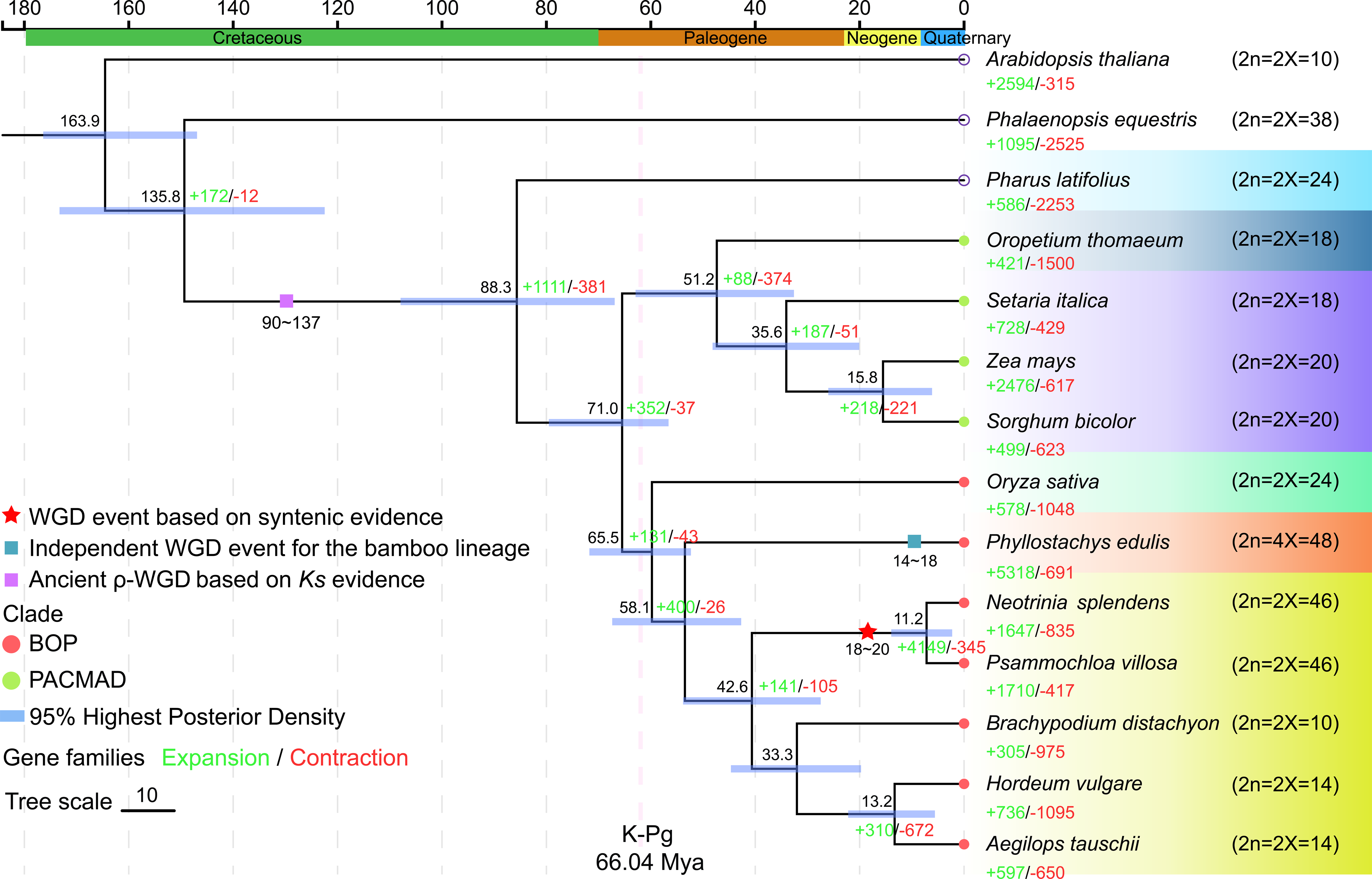
Phylogenetic and evolutionary analyses of *P. villosa*.

3.4 ***Whole genome duplication (WGD) and karyotype evolution of Psammochloa villosa*** The analysis of inter-chromosomal collinearity in *P. villosa* revealed the early evolutionary history of the genome included the occurrence of different rounds of WGD and/or tandem duplication. Intragenomic collinear analysis using MCScanX based on all-against-all identified 32,347 collinear gene pairs in 564 homologous blocks of the *P. villosa* genome. As shown in the dot-plot of syntenic segments, the collinear region located in chromosomes 1 and 2 of *P. villosa* was the largest homologous block, with 1,211 gene pairs, plus well-preserved syntenic region between chromosomes 8 and 10, 9 and 11, 12, and 16, 13 and 14, 22 and 23, supporting the occurrence of one or more large-scale duplications in the evolutionary history of *P. villosa* (Figure S7). Given that the annotated genome sequences of both *B. distachyon* and *O. sativa* were currently available, using the same criterion, we found 141 and 172 homologous blocks in the genomes of *O. sativa and B. distachyon*, including 5790 and 5052 collinear gene pairs (Figure 3), respectively. High intra-genomic gene collinearity provided evidence for the existence of two sub-genomes of *P. villosa*.

**Figure 3.**
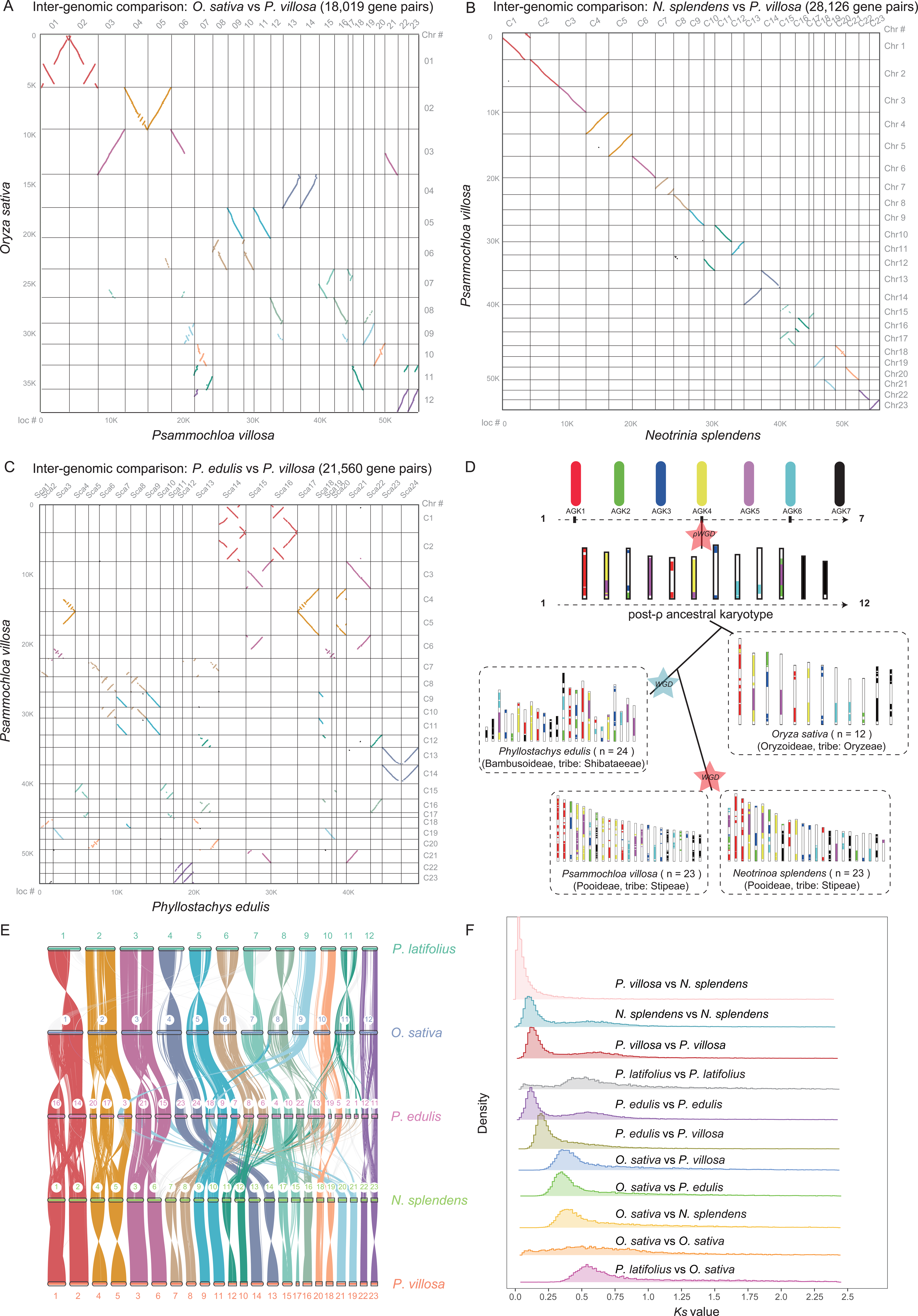
Identification of genomic syntenic blocks and protochromosomes 1-7 within the ancestral karyotype of grass (AGK).

The distribution of synonymous substitutions rate (*Ks*) of paralogs within *P. villosa* exhibited a bimodal peak at *Ks* values of 0.182 and 0.797, suggesting the occurrence of two WGD events (Figure S8a). In contrast, only a clear peak at ∼0.8 was evident in the *Ks* distributions for *O. sativa* and *B. distachyon* paralogs, pointing to a possible common WGD event (cWGD) at ∼96 Mya. Integrating this calibrated time, the recently WGD event shared between *P. villosa* and *N. splendens* was inferred to have occurred around 13-19 Mya (Figure S8b). The orthologous peaks for *P. villosa*, *N. splendens*, and *O. sativa*, corresponding to *Ks* values of ∼0.0971 and 0.43, were dated to 2.573 and 64.02 Mya, respectively, consistent with phylogenetic results. Comparing the *P. villosa Ks*-based WGD with orthologs between *P. villosa* and *N. splendens* (or *P. villosa* and *O. sativa*) genomes showed that this WGD predated the segregation between *P. villosa* and *N. splendens* but was later than the divergence between *P. villosa* and *O. sativa*. Therefore, the analysis suggests that a round of recent WGD event shared in the lineage of *P. villosa* and *N. splendens* genome, earlier than their divergence event (Figure S8c).

For inter-genomic gene collinearity, we detected 1,553 homologous blocks containing 35,753 collinear gene pairs by mapping *O. sativa* onto the *P. villosa* genome to determine orthologs after WGD events. If there was no gene loss, we anticipated that each collinear gene of *O. sativa* (or chromosomal area) would match genes in *P. villosa* for an orthologous and a paralogous gene (or chromosomal area). However, in genomic synteny analysis and dotplots, we observed that almost every collinear block from rice had two best-matched chromosomal regions in *P. villosa*. Additionally, we constructed a micro-synteny between *O. sativa* chromosomes (2n = 2x = 24, Oschr10 and Oschr11) and *P. villosa*’s pseudochromosomes 7, 18, 20, indicating that the ancestor of Os11 may have split into two segments. In the evolutionary history of *P. villosa*, these two segments joined together to form a larger region, which fusion resulted in the reduction from two sets of twelve duplicated pseudochromosomes produced by WGD to 23 chromosomes (Figure 3e, Table S14).

To further explore the chromosomal reorganization and karyotype evolution in *P. villosa* after the WGD event, we described the ancestral grass karyotype (AGK) and analyzed genome synteny (Figure 3d). We found genomic collinearity blocks on the chromosomes of five closely related species (*N. splendens*, *B. distachyon*, *O. sativa*, *P. villosa*, and *H. vulgare*) and observed that *P. villosa* has 18,852 AGK genes, accounting for 17.55% of all genes, lower than *B. distachyon* (47.79%), *O. sativa* (55.76%), and *N. splendens* (30.48%) but closer to *H. vulgare* (16.71%). We found that the section from eight chromosomes (3, 4, 5, 6, 7, 11, 13, 14) in *P. villosa* inherited more than two ancient chromosomes from AGK genes, which made the makeup of each chromosome more complicated than in the other four species. In particular, all five species had at least one evolutionarily conserved chromosome, and almost all AGK genes originated exclusively from ancient chromosome 1. For example, *P. villosa* had AGK genes on chromosomes 1, 2, and 9, while *N. splendens* had them on chromosomes 2, 3, 10, and 15 (Figure 3e).

### 3.5 The expanded PvXTHs gene families contributed to drought tolerance

Obviously, *P. villosa* and *N. splendens* underwent an additional WGD shared within Stipeae lineages, resulting in significant expansion and contraction of massive gene families. Xyloglucan endotransglucosylase/hydrolase (*XTHs*) gene family in *P. villosa* was taken as a case study due to its unique function in catalyzing xyloglucan hydrolase, which played an essential role in guard cell differentiation and stress responses (Fu et al., 2019; Rose et al., 2002). We found that 76 *PvXTH* genes had expanded in *P. villosa* (Table S15). Among these expanded genes, there were only 59 genes with both Glycosyl hydrolases-family 16 and Xyloglucan endo-transglycosylase C-terminus domains in *P. villosa*, which were distributed across all chromosomes (Figure S10). We confirmed the conserved domains and isoforms, leading to 59 *XTHs* genes being matched to BOP cluster. The 59 *XTHs* genes were then classified into three subgroups using a phylogenetic tree (Figure S11). This families’ peptides have two characteristic conserved domains (DEIDFEFLG), which were considered catalytic sites for hydrolase and transferase activity. Only two XET_C domains (PvT39296.1 and PvT39297.1) and one *XTH16* domain (PvT300781.1) contained an adenine at position 35 (Figure S12). We further identified conserved ExDxE domains, N-linked glycosylation sites, and conserved secondary structural loops 1-3 in the encoded proteins of these individuals through a conserved motif analysis (motif 2, Figure S13). Multiple alignments of 21 peptides from the nearest species revealed six conserved motifs, with the recognized trait (A/S substitution) of plants found at position 5 in motif 2 (Figure S14). Investigation of cis-acting elements in *XTHs* promoter regions indicated several cis-acting elements involved in drought responsiveness and responses to multiple hormones, such as the MBS element and typical regulatory elements like CAAT boxes and TATA boxes (Table S16, CAACTG). Under 20% PEG-6000 stress, the low level of Pv*XTHs* expression may protect plants by altering the cytoarchitecture (Figure S15). Pv*XTHs* genes showed strong collinearity (Figure S16) and purifying selection, as evidenced by the *Ka*/*Ks* values ≤ 1 for all homologous gene pairs (Table S17).

### 3.6 Population structure, genetic divergence and demographic history

To investigate the population genetic structure of *P*. *villosa*, 126 individuals from 42 sampling locations were used, representing the entire distributions range in China (Figure 4a). After strict filtering, a total of 1.87 Tb of genome resequencing data was generated, resulting in 8,774,438 high-quality SNPs and 1,326,289 InDels, respectively. Based on the high-quality SNPs dataset, population genetic analyses revealed three distinct lineages, which were geographically and climatically separated groups as inferred from PCA, ADMIXTURE and the phylogenomic tree. The PCA plot separated the three lineages along the first two eigenvectors (PC1 and PC2), which explained 33.65% and 9.91% of the total genetic variation, respectively. The third eigenvector (PC3) explained an additional 5.21% of the variation (Figure 4b). Phylogenetic analyses also supported the intraspecific division into three genetic clusters, and admixture analysis with *K* = 3 further corroborated this division, with the clusters henceforth referred to as northwest, northeast Helan mountains, and northcentral lineages (Figure 4c). Linkage disequilibrium (LD, measured as *r^2^*) declined with increasing physical distance in all three lineages. The northwest and north-central lineages reached half of their initial *r^2^* levels over shorter distances, whereas the northeast lineage showed a longer LD half-decay distance and consistently higher *r^2^* values, indicating more extended LD in this lineage (Figure 4d). This difference reflects contrasting genomic linkage backgrounds among the three lineages.

**Figure 4.**
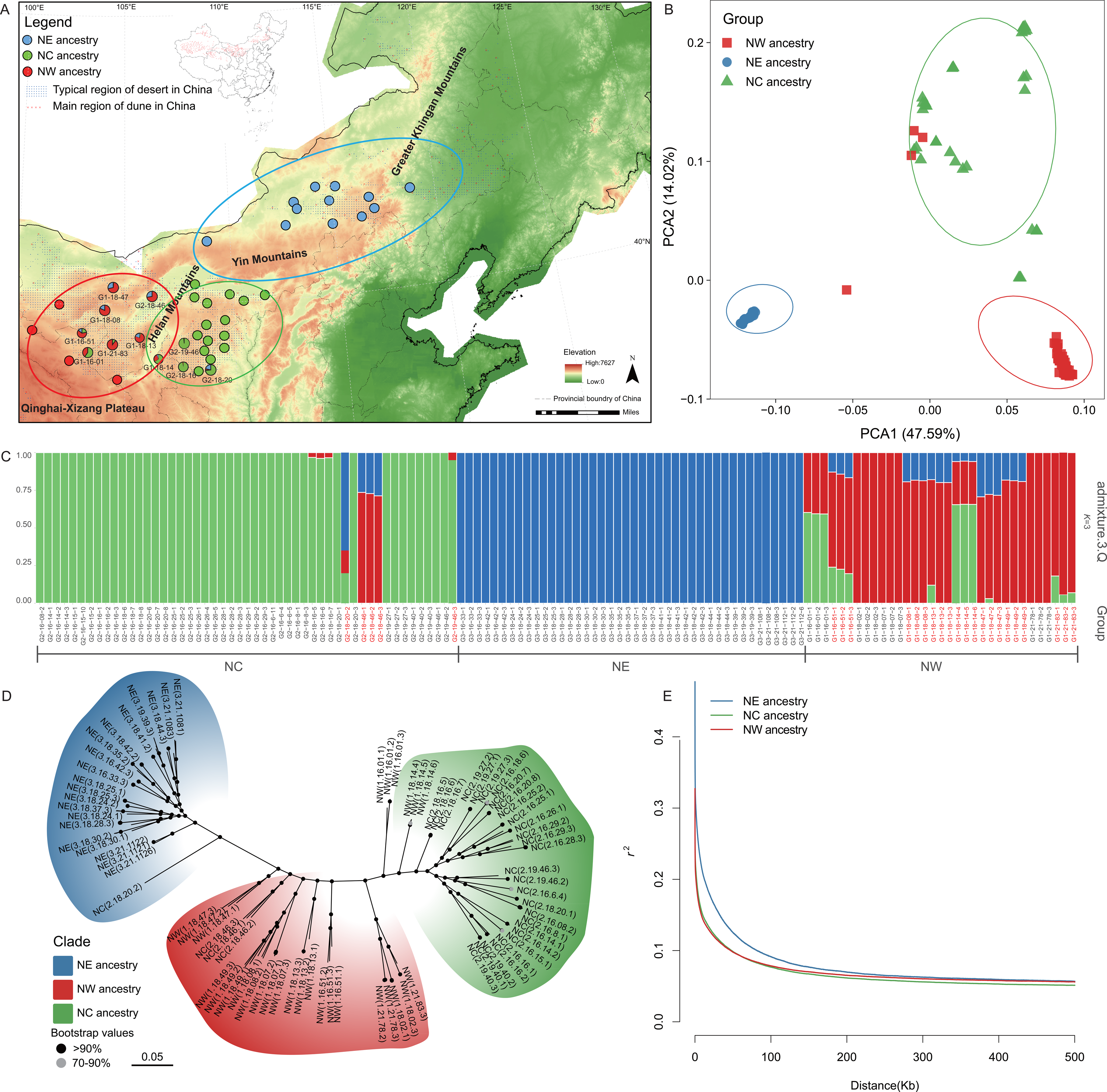
Population genomics analysis of *P*. *villosa*.

Because ancestry admixture can confound assessments of genetic diversity, population history and selection, we conduct calculations of genome-wide divergence (*F_ST_*) and the nucleotide diversity (*θπ*) relies on decreased population diversity to identify selection regions. Using 20kb sliding windows, genome-wide *F_ST_* scans revealed heterogeneous levels of genetic differentiation across the 23 chromosomes among the three lineage comparisons, and highly differentiated windows were subsequently evaluated together with lineage-specific reductions in nucleotide diversity to identify candidate selective regions. Based on the empirical top 5% thresholds, the joint analysis of *F_ST_* and *θπ* identified genomic windows showing both elevated population differentiation and lineage-specific reductions in diversity, which were considered candidate selective regions (Figure 6a). We also found that the northwest groups with mean Tajima’s *D* values = 1.162 was lower than northeast with 1.788 using 20 kb window sweep for each lineage, which group had 132,373 SNP sites with Tajima’s *D* values < 0. Examination of the candidate regions revealed local reductions in Tajima’s *D*, including negative values in several regions, which spatially coincided with elevated *F_ST_*, reduced *θπ*, and/or elevated XP-CLR scores (Figure 6b). Cross-population composite likelihood ratio (XP-CLR) scans further identified genomic regions with significant multilocus allele-frequency differentiation among the three lineage comparisons, with 1,301, 611, and 984 windows remaining significant after FDR correction (FDR < 0.05) for the northeast VS north-central, northwest VS north-central, and northwest VS northeast comparisons, respectively (Figure S19a-c). These XP-CLR signals were further compared with the *F_ST_*, *θπ*, and Tajima’s *D* profiles across representative candidate regions (Figure 6b).

Finally, a gene-wise HKA-based test identified 71 genes showing significant departures from the genome-wide ratio of fixed to polymorphic sites (*P* ≤ 0.05) in all three lineages (Table S20). To obtain a more stringent set of candidate selective regions, the genomic windows identified by the joint *F_ST_*-*θπ* scan were intersected with significant XP-CLR signals. Of the 738, 664, and 994 candidate windows initially identified for the northeast VS northwest, northeast VS north-central, and northwest VS north-central comparisons, respectively, 162 (22.0%), 116 (17.5%), and 247 (24.8%) showed genomics overlap with XP-CLR signals. These concordant windows were therefore retained as candidate selective regions supported by both genetic differentiation/diversity reduction and cross-population allele-frequency differentiation (Table S21). Therefore, we have screen gene under local selection by combining *F_st_* and *θπ* and identified 2375 select region (including 509 genes), in which 50 *PvXTHs* gene families had undergoing strongly selection between northwest and northeast lineages of *P. villosa* (Figure 5, Table S18).

**Figure 5.**
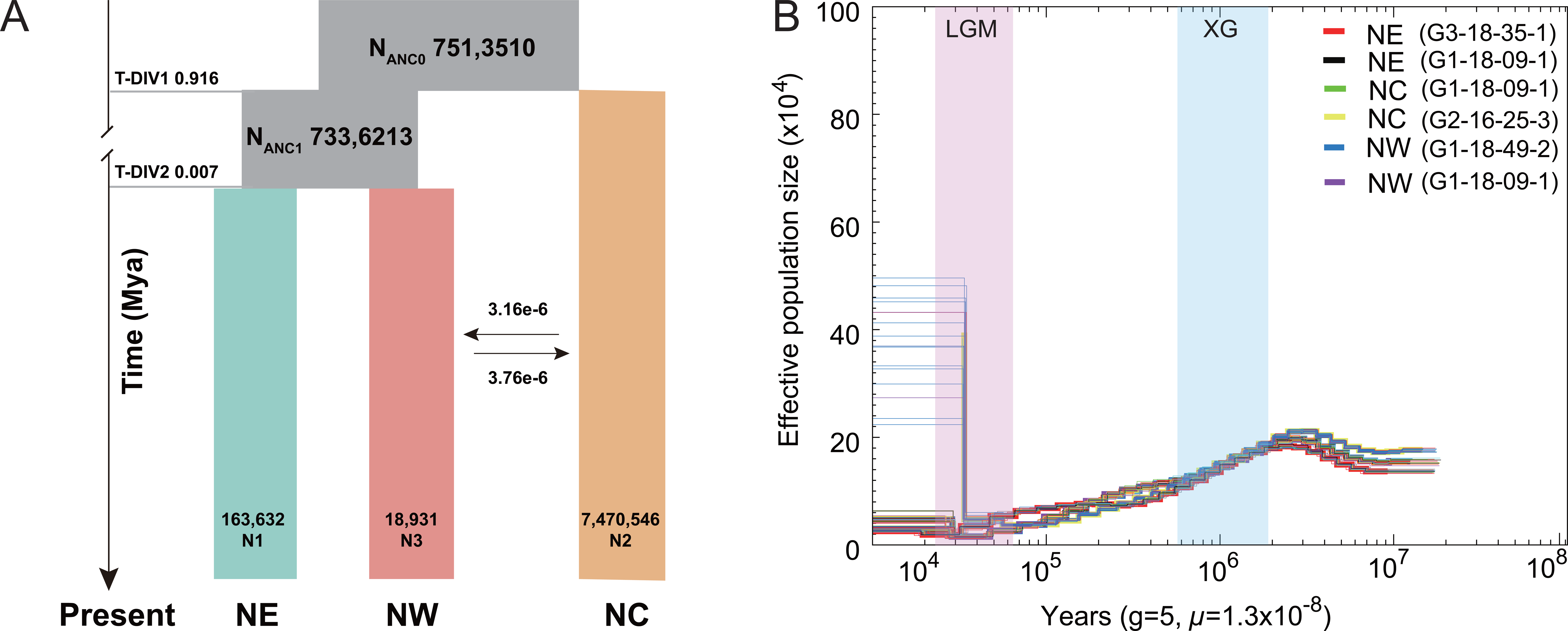
Demographic history of three lineages of three lineages of *P*. *villosa*.

**Figure 6.**
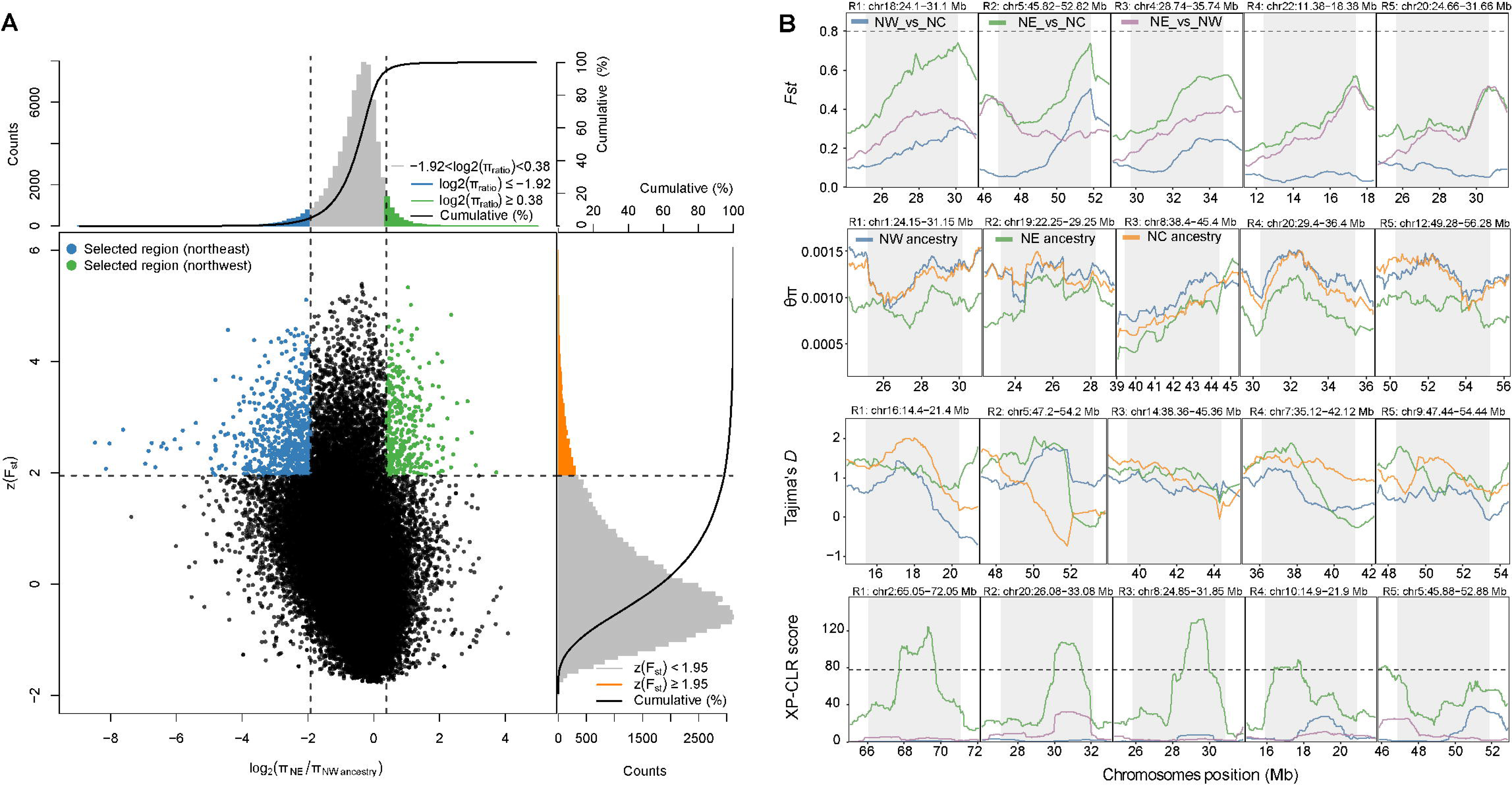
Genomic regions with strong selective sweep signals in the lineages of *P*. *villosa*.

Demographic histories and temporal changes in effective population size (*Ne*) of the northeast, northwest, and north-central lineages were reconstructed using PSMC (Figure 5b) and SMC++ (Figure S15), with broadly concordant demographic trajectories obtained from the two approaches. The three lineages showed a pronounced increase in *Ne* beginning at approximately 8.0 Ma, followed by a decline toward approximately 0.3-0.1 Ma and modest subsequent increases toward the recent past (Figure 5b).To further characterize the demographic relationships among the three lineages, alternative demographic models were evaluated using fastsimcoal2. The best-supported model inferred two divergence events at approximately 0.916 and 0.007 Ma. The ancestral effective population sizes were estimated at approximately 7.51 × 10^6^ and 7.34 × 10^6^, whereas the current effective population sizes were 163,632, 18,931, and 7.47 × 10^6^ for the northeast, northwest, and north-central lineages, respectively. The model also inferred bidirectional migration between the northwest and north-central lineages, with estimated migration rates of 3.16 × 10^-6^ and 3.76 × 10^-6^ in the two directions (Figure 5a). Overall, these demographic estimates were broadly concordant with the reconstructed population-size trajectories of the three lineages.

Gene flow in the recent past was much higher than in the more distant past. Furthermore, gene flow in different divergence stages was asymmetrical, with a higher rate of gene flow from northwest to northcentral lineages (Figure S21).

### 3.7 Identification of genomic variants associated with ecological adaptation

The high-quality reference genome for *P. villosa* coupled with the resequencing data facilitated the precise characterization of genomic information including SNPs and Indels. We used two complementary approaches to identify genetic variants associated 46 environment and climate factors. Of the 324,223 significant variants identified by LFMM, we identified a total of 6341 protein-coding genes that were significantly related to one or more factor (Figure S22). Although LFMM approaches are univariate, testing for associations between one variant and one environmental factor at a time, we still detected PvG00000047987, a gene belonging to the *PvXTHs* gene families, as associated with precipitation and wind (Figure 6). Furthermore, after considering the ranked importance using the gradient forest analysis and the multicollinearity of these correlated variable, four variables with Spearman correlation coefficient |*r*|< 0.6 were retained for the RDA analyses, including a temperature variable (Figure S23, mean temperature of warmest quarter (BIO10), precipitation seasonality (coefficient of variation, BIO15), maximum precipitation of warmest quarter (BIO18)) and vapor pressure of December (vapr12). RDA analysis reveal that climate-associated genetic variation of population was visualized by these four climatic variables. RDA1 and RDA2 axes account for 87.79% of the total variance in the data, and mean temperature of warmest quarter was negatively associated with both RDA1 and RDA2, highlighting their adaptive divergence within the population (Figure S24). We found that these variants largely overlapped within the significant variants by LFMM. Of 302 SNPs were found to display extreme loadings (standard deviation > 3) along one or multiple RDA axes (Table S19). These shared variants were regarded as “core adaptive variants” of *P. villosa*.

## Discussion

### 4.1 Comparative genomics reveals the evolutionary genome of P. villosa

The high-quality, chromosome-level genome assembly and annotation for *P. villosa*, a key and dominant species in desertified meadows in Inner Mongolia of China, provides unique resources for studying the genomic basis of xerophytic ecological-adaptation to desertification region. Based on our previous karyotype and chromosome ploidy analysis from different *P. villosa* populations (Wang et al., 2021), we conducted a genome survey and *K-mer* analysis, revealing the comparatively high repetition (75.38%) and heterozygosity ratio (1.15%) genome. The results were even more complex than for the closely related species *N. splendens*, which had 62.11% and 1.13%, respectively (Ren et al., 2022). These characteristics make the genome of the lineage of Stipeae tribe (typified by *P. villosa* and *N. splendens*) very complex and challenging to analyze and interpret. Phylogenomic and synteny analyses indicate that *P. villosa* and *N. splendens* shared a whole-genome duplication 13–19 Ma and diverged ∼7.2 Ma, which is in coincident with Late Miocene interior Asia aridification driven by global cooling and Tibetan Plateau uplift (Liu et al., 2016). Undoubtedly, the unique physical and geological history of the arid regions in Northwest China have profoundly influenced plant species distribution and evolution (Zheng et al., 1992; Harrison et al., 1998; Guo et al., 1999). These long-term selection stresses, such as drought, high temperatures, frequent strong winds, and extensive habitat fragmentation and desertification caused by Quaternary climate changes and the uplift of the Himalayan-Tibetan Plateau have driven plant populations to undergo functional gene subfunctionalization (Hu et al., 2023; Wang et al., 2025), adaptive evolution (Ma et al., 2013), and species diversification (Muellner, 2019; Wang et al., 2023). The unique gene families of *P. villosa* were enriched in "guard cell differentiation", "response to oxidative stress, water", "peroxidase activity" and "xylosyltransferase activity" Gene Ontology (GO) categories. Based on sequence homology, these unique gene families of *N. splendens* were predicted to "response to oxidative stress" pathway. We also identified gene duplication originating from their expend gene families both have multiple isoforms, including but not limited to *ABI1*/*ABI2*, *BIN2*, *BES2*, *SnRK2.2*/*SnRK2.4*, *PYL10*/*PYL12* (Table S13). A high-quality genome facilitates the accurate description of new germplasm and enables the investigation of the genetic basis of drought tolerance. These results also provide crucial insights into the functional organization and potential evolutionary history of the *P. villosa*’s pangenome, with implications for a wide range of research fields.

Obviously, frequent activation of repetitive elements is one of the primary driving forces that increase genome sizes (Flavell et al., 1974; Levin & Moran, 2011). LTR-RTs are the most prevalent of repetitive elements in plants (Vicient et al., 1999). In the *P. villosa* genome, most LTR-RTs have expanded and accumulated recently (Figure S3), indicating that *P. villosa* has undergone a recent burst of TE activity. This could potentially impact its genomic stability, gene expression regulation, and evolution (Wendel, 2000). We identified a total of 54,317 protein-coding genes in the assembled genome of *P. villosa*. This number is greater than those in other Poaceae species included in our analyses, such as *O. sativa* and *B. distachyon*, which have not undergone an additional WGD. Our finding implies *P. villosa* exhibits a high degree of genetic complexity and variation, which may underlie various adaptations and evolutionary innovations that have enabled *P. villosa* to thrive in its natural environment.

### 4.2 Chromosomal variation and reduplicated genes retention post-WGD provide new insights into the plant’s drought adaptability

Ancient whole-genome duplications are widespread in plants and are strongly supported by evidence found in species-rich lineages of angiosperms (Doyle et al., 2008; Escudero & Wendel, 2020; Jiao et al., 2011; Ren et al., 2018; Wu & Jiao, 2020). The Pooideae subfamily, the largest grass subfamily, encompasses over 3,900 species. Zhang et al. (2022) suggested that four-fifths of Pooideae species shared a cluster of gene duplications during the Eocene-Oligocene transition, coinciding with elevated diversification rates in this lineage. The primary contribution of WGD events to species genomic variation is primarily through providing new genetic material, accelerating mutation and selection processes (Hämälä et al., 2024). Several empirical studies have demonstrated that the retention of gene duplicates after WGD depends on their function, and the proportion of retained duplicates typically decreases rapidly over time (Ren et al., 2018). This retention increases the sizes of gene families and creates opportunities for duplicate copies to participate in lineage-specific adaptive changes (McDaniel, 2024). Although the role of adaptive forces in establishing polyploids remains a subject of debate (Murat et al., 2010; Mayrose et al., 2011; Soltis et al., 2014), we hypothesize that prior to the divergence of closely related species within the Stipeae tribe, they likely underwent a shared initial polyploidization event (Tkach *et al*., 2021). Our genomic collinearity analysis of *P. villosa* underwent an additional WGD event estimated to have occurred 13-19 Mya (Figure S9) in the stem node of the clade of *P. villosa* and *N. splendens* (Subfamily: Pooideae, Tribe: Stipeae). We identified a later divergence event between *P*. *villosa* and *N. splendens*, a finding that aligns with the ‘lag-time model’, which posits that diversification typically follows whole-genome duplication (WGD) events after a lag period (Schranz et al., 2012). Therefore, we focused on duplicate genes that increased gene copy numbers and facilitated functional innovation in *P. villosa* and *N. splendens* compared to other grass species. It is not surprising that 48 expanded gene families in the genome of *P. villosa* were comparable to those of species that diverged subsequent to this recent WGD event, paralleling the duplication pattern in the *N. splendens* genome, in which each *O*. *sativa* gene locus (2n = 2X =12) corresponds to two best-matched orthologous, and two outparalogous genes, which, based on a phylogenetic tree, form three subgroups most closely related to *AsXTHs* (Figure S11). Widespread WGDs were correlated with the emergence of lineages with high degrees of diversification and periods of global climate change (Xu et al., 2011). Importantly, our study highlights the retention of reduplicated genes after WGD events in driving plant lineages evolution and diversification. This finding supports the lineage-specific WGD event is likely to have played a crucial role in species diversification within the Stipeae tribe.

### 4.3 Genomic islands of divergence and ecological adaptation

Previous research on *P*. *villosa*, analyzing 1,728 AFLP markers, has shown plentiful genetic variation primarily within populations but infrequent gene flow (Lv et al., 2021). The geographic isolation caused by Yinshan-Helan Mountains and large deserts area (e.g., Tengger Desert; Mu Us Sandy Land) may limit long-distance dispersals events. While corroborating previous studies, our SNP-based analyses resolve the population into three primary lineages. The NE ancestry had slower the rate of LD decay and lower level of mixed-lineage (Figure 4d). This suggests that the NE lineage was older and exhibited a weaker genotype correlation among individuals based on Hardy-Weinberg equilibrium. In contrast, the NW and NC lineages exhibit more rapid LD decay, reflecting their closer genetic relationship and higher levels of inter-lineage gene flow (Figure 4c). We also found that the level of intraspecific divergence was heterogeneous and genomic islands of divergence were scattered across the genome. Similar to the genomic divergence pattern observed under the speciation-with-gene-flow model (Feder & Nosil, 2010; Hu et al., 2021; Ma et al., 2018). For example, we calculated genetic divergence (*F_st_*) between the NE and NW lineages, and the results were significantly higher than those of other inter-lineages. Additionally, the NC lineage exhibited higher polymorphism with a mean *θπ* = 0.0013 compared to NW lineage. After analyzing gene flow by TreeMix, we found a single migration from NC to NW, which aligned with the observed population structure (Figure S22). Therefore, it is more reasonable to classify the population’s phylogeographic structure in *P*. *villosa* into three subgroups.

We sought to estimate the paleogeographic and climatic drivers of demographic history for *P*. *villosa*, which features a modest increase in *Ne* since ∼10 Ma followed by a continuous decline from ∼0.1 Ma. Previous studies have shown that Late Miocene (∼8 Ma) global cooling together with Qinghai-Xizang Plateau (QXP) uplift drove aridification of the Asian interior (Liu et al., 2016; Murat et al., 2010). Using demographic history reconstructions, we find that the *Ne* of *P. villosa* lineages has steadily declined since the Late Miocene, coinciding with regional desert expansion. Given the sustained *Ne* contraction since the Late Miocene, we anticipated that the relative strength of genetic drift and nature selection had been increased, generating heterogeneous patterns of divergence that manifest as genomic islands detectable by selection scans. Genetic differentiation and polymorphism of different genes should be consistent in the absence of selection based on neutral theory (Lewontin & Krakauer, 1973). Although previous studies highlighted the genetic variation underlying intra-population, the contribution of gene flow in generating the genomic islands could not be excluded here. We conducted the other approach based on difference of site frequency spectrum (SFS) for multi-locus allele (Yang et al., 2023), such as XP-CLR and HKA test, to identify selection sweeps between NE and NW lineages (Table S20). Our results showed strongly selection signals between NE and NW lineages (Figure S20a). These results may suggest the occurrence of divergence hitchhiking or selection within regions, both of which tend to increase *F_st_* or LDdecay distance and reduce genetic diversity, and thus also contribute to the formation of genomic islands.

The “core adaptive variants” of the *P. villosa* lineages to drought environment were identified by LFMM and RDA analysis. In several candidate loci associated with 43 climatic variates and geographical factors (Longitude latitude altitude), *PvXTHs* (PvG00000047987) was significantly related to wind and precipitation factor (Figure 6). The result was in accord with the selected region of inter-lineages (between NW and NE lineages) via combining *θπ* and *F_st_* (Figure 5). Previous research (Sang et al., 2022) reported that the variants of *CRL1* gene associated with precipitation displayed similar geographic distribution in frequencies, leading to the *Populus koreana* adaptation to local environment. Our results suggested that these differentiation alleles associated with climatic change, specifically in functional genes, have dominated lineage local adaption though multiple pathways. Not only *PvXTHs* observed difference of *Ks* in the exon region, the geographic distribution of its allele frequencies showed differentiation. It is plausible that local adaption of local selection has contributed to the genetic divergence of xerophytic ecological-adaptation to desertification regions. However, due to the deficiency to explore the genetic transformation system for this xerophyte perennials species, we have attempted to identify stable reference gene using qRT-PCR and the integrated statistic method (Liu et al., 2022b). In future, investigation on these core adaptive gene’s functions and verification experiment will profile its expression under abiotic stress. Our research provides a well-assembled genome and genome-level perspective for the genetic evolution and ecological adaption of *P. villosa*.

## Conclusion

We present a high-quality chromosome-scale genome assembly of *P. villosa*, a dominant species under water-deficient conditions, as a resource for studying drought tolerance of plants. Comparative analysis of this genome, combined with other available genomes, allowed us to infer the phylogenetic relationships, investigate species divergence, and the origin of *P. villosa*, and identify gene families involved in environmental adaptations. Our analyses revealed a bimodal pattern in the paralogous peaks in *P. villosa*, indicating two rounds of whole-genome duplication (WGD) events. Following this, the genomes evolved separately between *N. splendens* and *P. villosa* and from the ancestral grass karyotype. Therefore, we have identified a chromosomal rearrangement which was involved in stress resistance and metabolic pathways. Relatively, intraspecies genetic differentiation unmasked the population structure and gene flow barriers of *P. villosa*. We identified 2,375 selected regions (including 509 genes), and annotated 50 *PvXTHs* genes undergoing strongly selection. In addition, precipitation seasonality and three other variables were of most importance in contributing to intraspecific variation, suggests that these fixed mutations may have played a crucial role in the dryland ecological adaptation and speciation of *P. villosa*. Our analysis of genotype-environment association and the high-quality reference genomes of *P. villosa* provides a valuable resource for further evolutionary genomics studies in this monotypic genus.

## Acknowledgements

This work was financially supported by the National Natural Science Foundation of China (No. 32160297, 41761009) and the Program of Science and Technology International Cooperation Project of Qinghai Province (No. 2023-HZ-810) made to Yuping Liu.

## Data availability statement

Genome assembly of *P. villosa*, Iso-seq and Hi-C data have been submitted to the China National Center for Bioinformation under BioProject number PRJCA050660: genome assembly and annotation have been deposited in the Genome Warehouse in National Genomics Data Center in accession number GWHHKIF00000000.1; Genbank databases (NCBI) for PacBio (HiFi) data and Hi-C data under : PRJNA1413959, with accociaiton number SRR37011173 and SRR37240158; Meanwhile, all of resequencing data to NCBI under accociation number SUB1598134; Iso-seq data–SRR11787985 and SRR11787359; We have submitted the RNA-seq data to NCBI under accociation number SRX10895848, SRX10895849, SRX10895850, SRX10895851, SRX10895852, SRX10895853, SRX10895854, SRX10895855, SRX10895847.

## Author Contributions

We greatly appreciated Ting Lv, Dandan Su and Yu Zhang leading of quantitative real time polymerase experiments, Ping Yang performing the germination experiment in the control chamber, Xiayu Hu assisting DNA, RNA extraction experiments, Changyuan Zheng and Gui Fu analyzing data of full-length transcriptome, and the anonymous reviewers for their helpful comments.

---

**Figure 7.** Manhattan plot for SNPs variants related to the climate factor.

**Table 1**. Classification of interspersed repeats in the assembled genome of *P. villosa*.

## Supplemental files

Figure S1. *K*-mer analysis revealed ploidy profiling of the genome in *P. villosa* by using *K*-mer = 17.

Figure S2. Hi-C assisted assembly of pseudochromosomes of *P. villosa*.

Figure S3. Insertion time of LTRs in four species, namely *P. villosa*, *O. sativa*, *B. distachyon*, *N. splendens*.

Figure S4. Comparison of gene length, mRNA length, exon length, intron length and CDS, exons number between *P. villosa* and eight related species.

Figure S5. Venn diagram of orthologous genes (gene families) shared among the seven species, *tauschii*, *N. splendens*, *B. distachyon*, *H. vulgare*, *O. sativa*, *P. villosa*, and *P. edulis*.

Figure S6. Go enrichment of the expanded gene families of *P. villosa*.

Figure S7. Overview of dotplots within the *P. villosa* genome for paralogous genes.

Figure S8. Collinearity analysis of interspecies genome based on frequency of synonymous substitution levels (*Ks* values and 4DTV) of syntenic orthologous.

Figure S9. A typical collinearity pattern between genomic regions of *P. villosa* and *N. splendens*.

Figure S10. Distribution of all *XTHs* genes on chromosomes of *P. villosa*.

Figure S11. The phylogenetic relationship of *XTHs* gene derived from *P. villosa, O. sativa*, *B. distachyon*, *H. vulgare*, and *N. splendens*.

Figure S12. Structural analysis of the ten conserved motifs in *XTHs* gene structures.

Figure S13. Sequence composition of conserved motif 2 of *P. villosa*.

Figure S14. Multiple alignment of motif 2 of *XTHs* for showing the conserved secondary structures.

Figure S15. Gene expression of the xyloglucan endo-transglucosylase/hydrolase family (59

*XTHs*) under different drought treatments using 10% (TM) and 20% (TH) PEG-6000. Figure S16. Genomics collinearity of the *PvXTHs* family (59 genes) by synteny of *P. villosa*.

Figure S17. Genetic diversity, population divergence between northwest and northeast lineages.

Figure S18. Genome-wide scanning to identify selection region by the reduction of diversity (ROD) northwest and northeast lineages.

Figure S19. The estimation of XP-CLR score among northwest, northeast and northcentral lineages.

Figure S20. Demographic history and effective population size of *P. villosa*.

Figure S21. Trees inferred by TreeMix. Gene flow test generated 126 independent simulations of 23 chromosomes from each population using the demographic model without migration, and inferred population trees.

Figure S22. Mapping the location of the 324,223 environmental adaptive genes detected by LFMM model across the whole genome.

Figure S23. The sort of accuracy importance for each variate’s contribution.

Figure S24. The redundancy analysis selected the four variables associated with population genetic variation.

Table S1. Geographic localities and population distribution of *P. villosa*.

Table S2. Estimation of the genome size of *P. villosa* based on 17-mer statistics.

Table S3. Summary of DNA sequencing data.

Table S4. Summary of assembly based on Hi-C data.

Table S5. Summary of Pacbio full-length cDNA sequencing

Table S6. Correction and statistics for genome of *P. villosa* by PacBio transcriptome data.

Table S7. Genome assembly completeness evaluation by BUSCO.

Table S8. LTR subclass ratio of *P. villosa* and other five genomes of Poaceae.

Table S9. Summary of predicted protein-coding genes annotation and their supporting evidences.

Table S10. Functional annotation of predicted genes in the genome of *P. villosa*.

Table S11. Structure annotation of predicted ncRNA in the genome of *P. villosa*.

Table S12. Summary of gene family clustering.

Table S13. Gene ontology (GO) enrichment analysis of the unique families in *P. villosa*.

Table S14. Collinearity of homologous between *O. sativa* and *P. villosa*.

Table S15. identification and confirm for *PvXTHs* gene families for *P. villosa*.

Table S16. The prediction of cis-element for *PvXTHs* gene.

Table S17. Duplicated events and tandem for *P. villosa XTH* genes.

Table S18. Genomic islands of divergence and *XTHs* annotation based on *F_st_* and *θπ* ratios between NW and NE lineages.

Table S19. The calculation variants associated with local environmental adaptation of by gradient forest in RDA analysis.

Table S20. Hudson-Kreitman-Aguadé (HKA) test for NC, NE and NW lineages of *P. villosa*.

