## Supplementary material for "Multi-Omics analysis provides crucial insights into ecological adaptation to dryland of a dominant grass (*Psammochloa villosa*, Poaceae) in Northwest China": Table 1

Table 1 Classification of interspersed repeats in the assembled *P. villosa* genome

| Class | Subfamily | Length(bp) | Percent (%) |
|---|---|---|---|
| Class I: Retrotransposon | Total LTR | 856,335,233 | 56.231 |
| | LTR/Copia | 260,793,668 | 17.125 |
| | LTR/Gypsy | 598,546,060 | 39.303 |
| | Other LTR | 11,687,619 | 0.767 |
| | SINEs | 965,104 | 0.063 |
| | Total LINEs | 41,713,334 | 2.739 |
| | LINE/L1 | 36,502,347 | 2.397 |
| | LINE/L2 | 3,314,654 | 0.218 |
| | Other LINE | 1,902,016 | 0.125 |
| Class II: DNA Transposon | Total DNA | 150,153,894 | 9.86 |
| | CMC-EnSpm | 92,330,059 | 6.063 |
| | hAT-Ac | 5,337,657 | 0.35 |
| | hAT-Tip100 | 1,593,869 | 0.105 |
| | PIF-Harbinger | 12,888,473 | 0.846 |
| | MULE-MuDR | 29,430,559 | 1.933 |
| | Other DNA | 10,093,247 | 0.663 |
| Unclassified | | 111,209,987 | 7.303 |
| Low_complexity | | 954,453 | 0.063 |
| Simple_repeat | | 48,550,279 | 3.188 |
| Satellite | | 2,816,043 | 0.185 |
| Total content | | 1,154,945,209 | 75.84 |
