## Supplementary material for "Multi-Omics analysis provides crucial insights into ecological adaptation to dryland of a dominant grass (*Psammochloa villosa*, Poaceae) in Northwest China": Figure S1-21

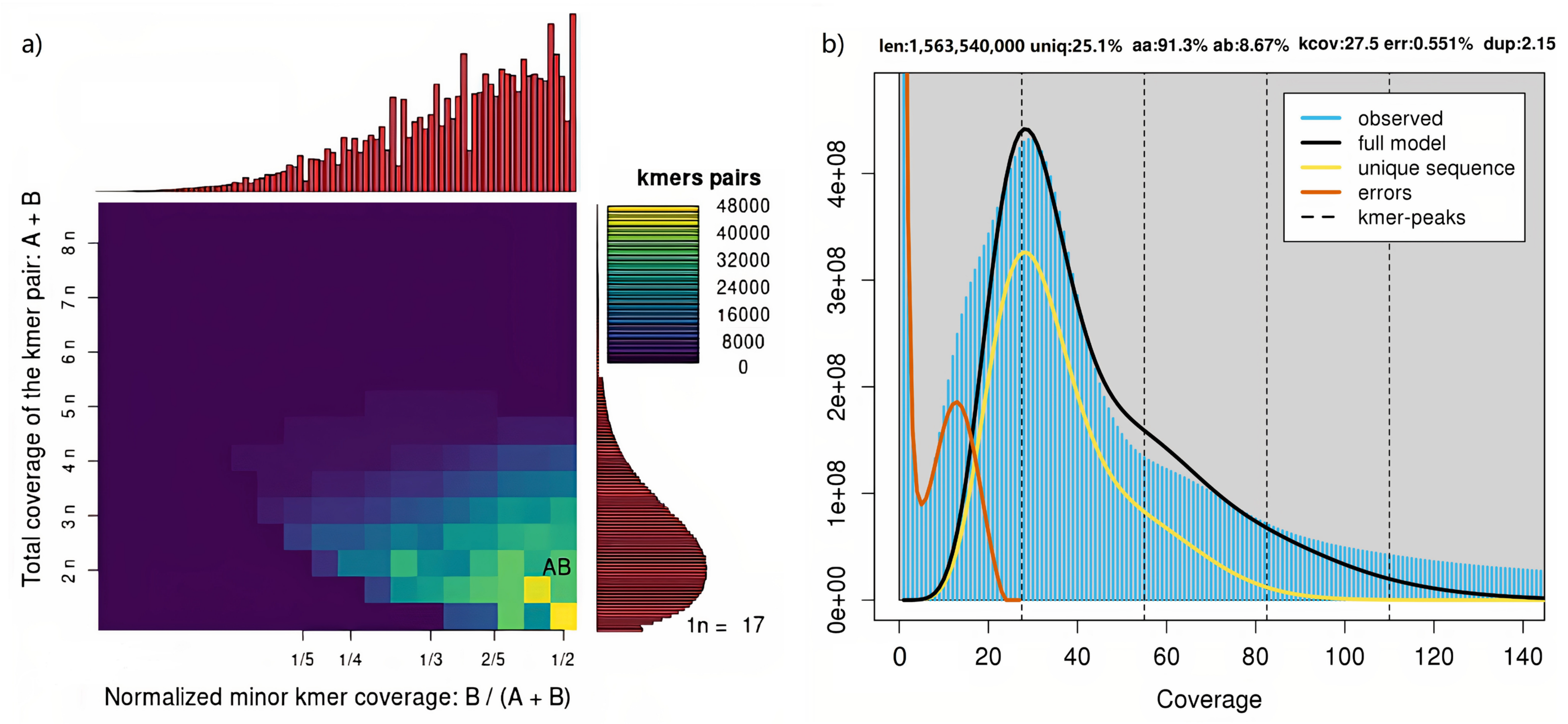


**Figure S1.** *K*-mer analysis revealed ploidy profiling of the *P. villosa* genome by using *K*-mer = 17. (a) The sharp peak depth was 13 in the distribution. There is not too obvious peak of hybrid peak and a very smooth peak (around 26) in *K*-mer frequency distribution curve (Kajitani et al., 2014). (b) The plot shows distinct "smudges" representing each *K*-mer pair with the greatest of density of *K*-mers relating to the ploidy level of the genome.

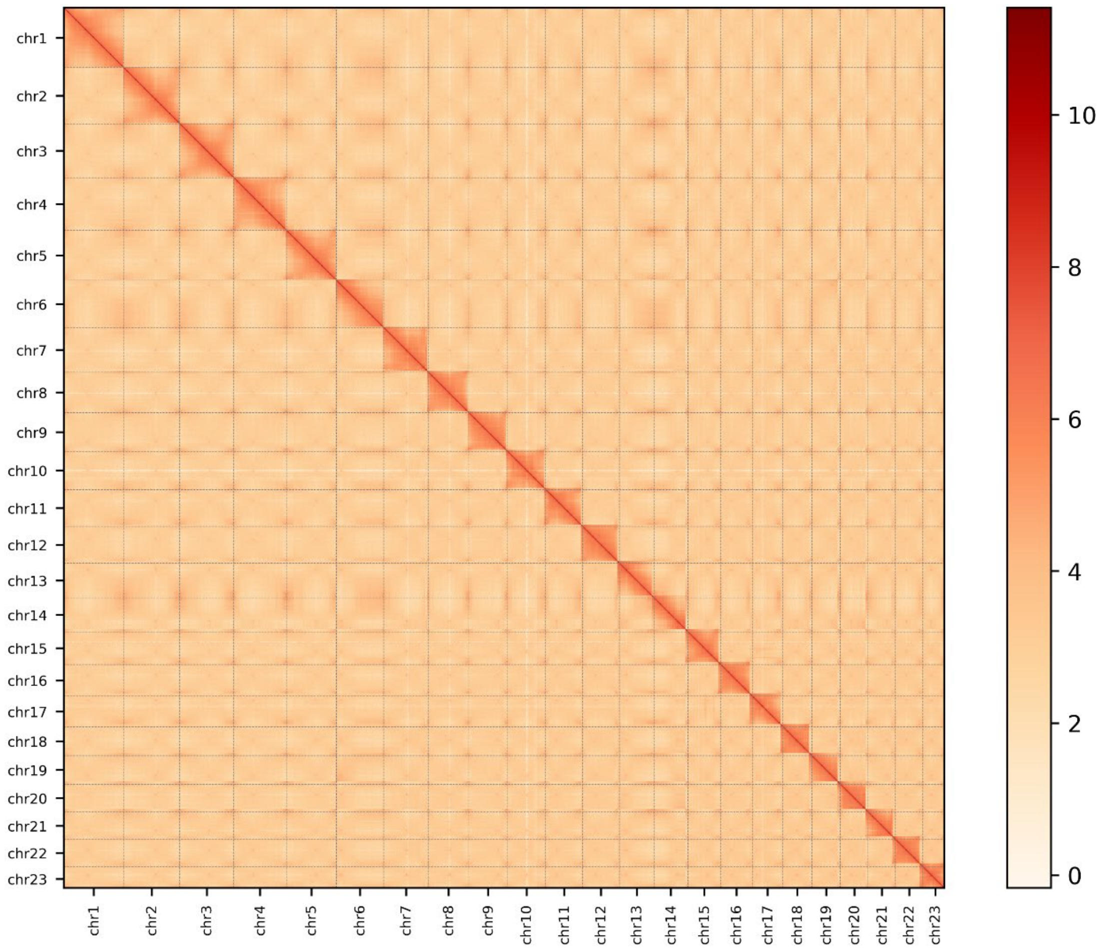


**Figure S2.** Hi-C assisted assembly of pseudochromosomes of *P. villosa*. Heatmap showing Hi-C interactions under a resolution of 200 kb, and the diagonal pattern for the intrachromosomal interactions may reflect the linear distance of chromatins.

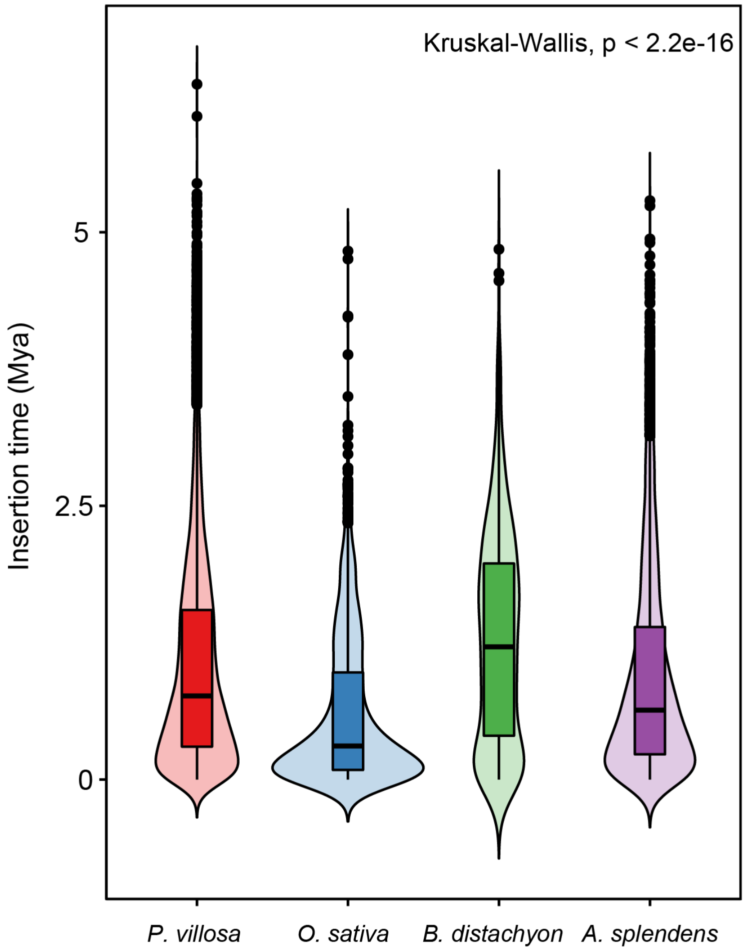


**Figure S3.** Insertion time of LTRs in four species, namely *P. villosa*, *O. sativa*, *B. distachyon*, *A. splendens*.

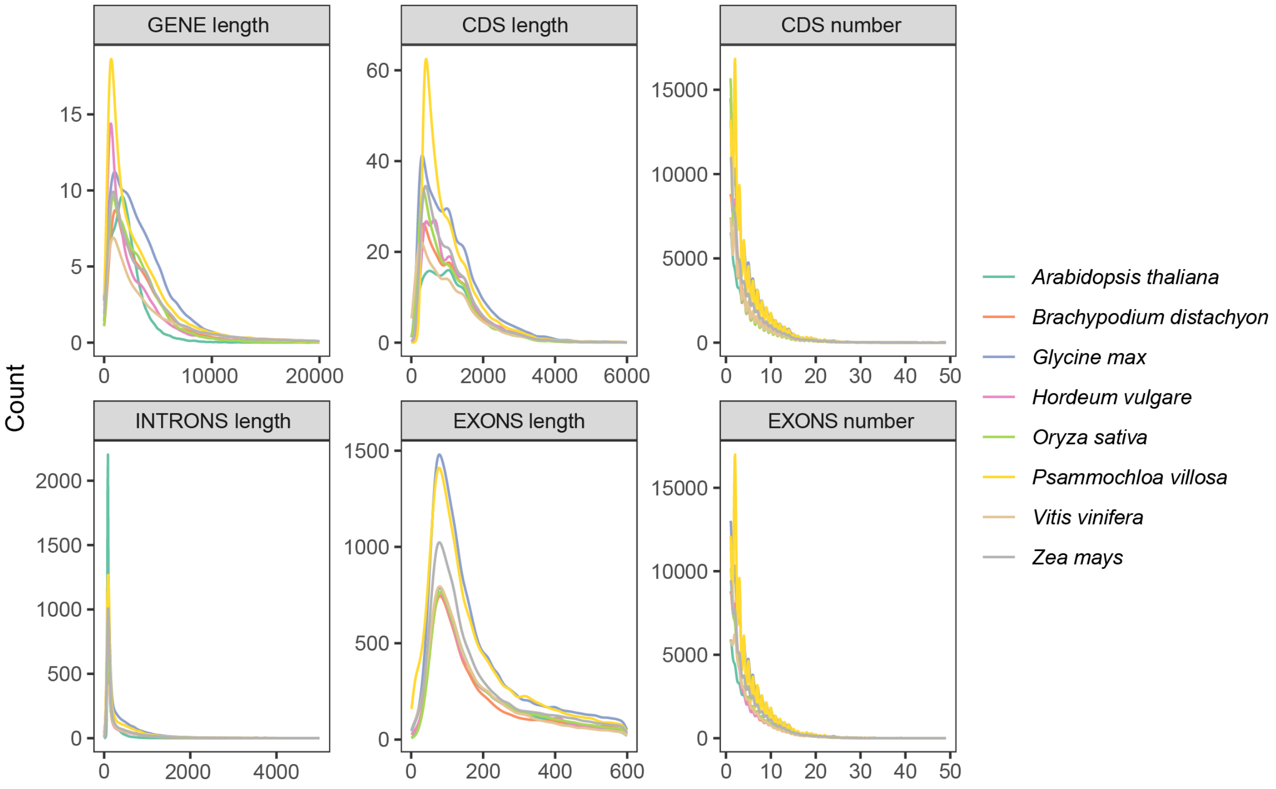


**Figure S4.** Comparison of gene length, mRNA length, exon length, intron length and CDS, exons number between *P. villosa* and eight related species.

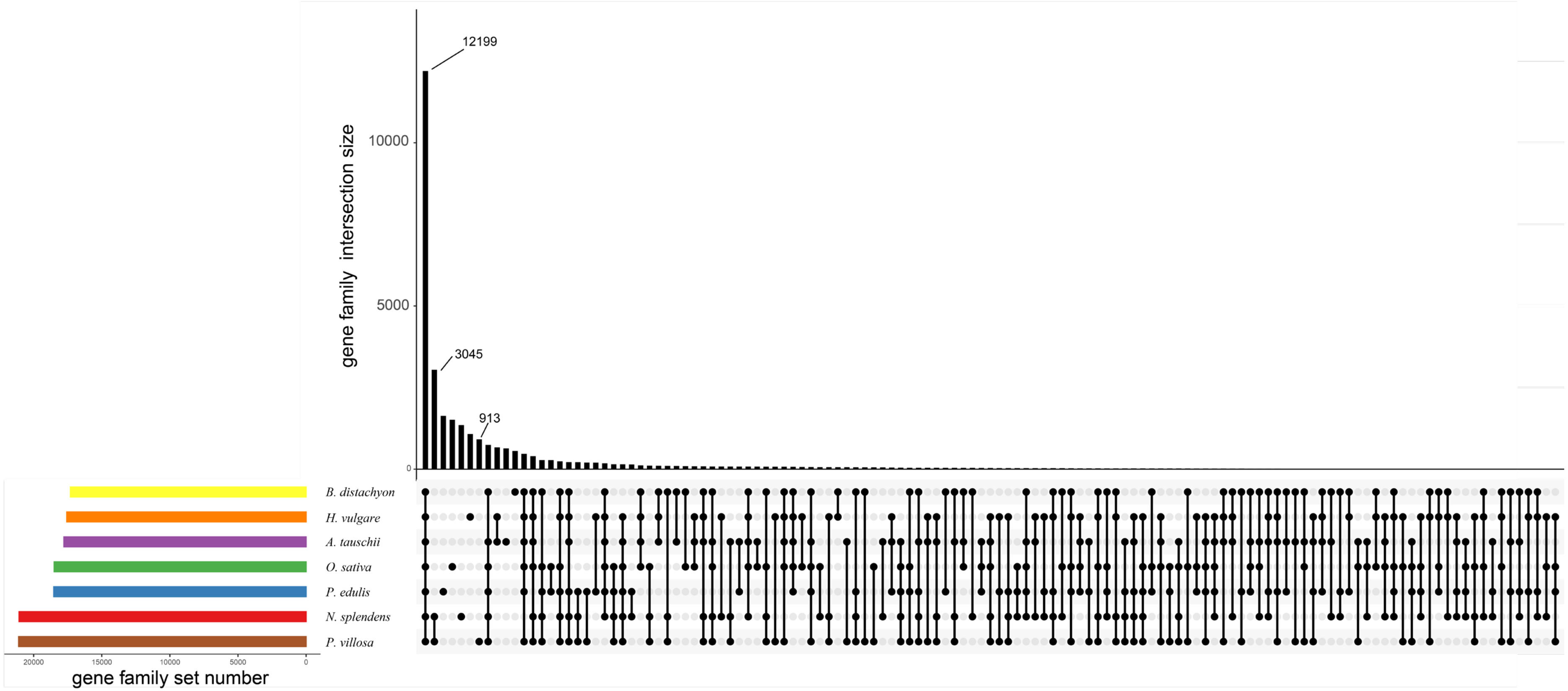


**Figure S5.** Venn diagram of orthologous genes (gene families) shared among the seven species, *A. tauschii*, *N. splendens*, *B. distachyon*, *H. vulgare*, *O. sativa*, *P. villosa*, and *P. edulis*.

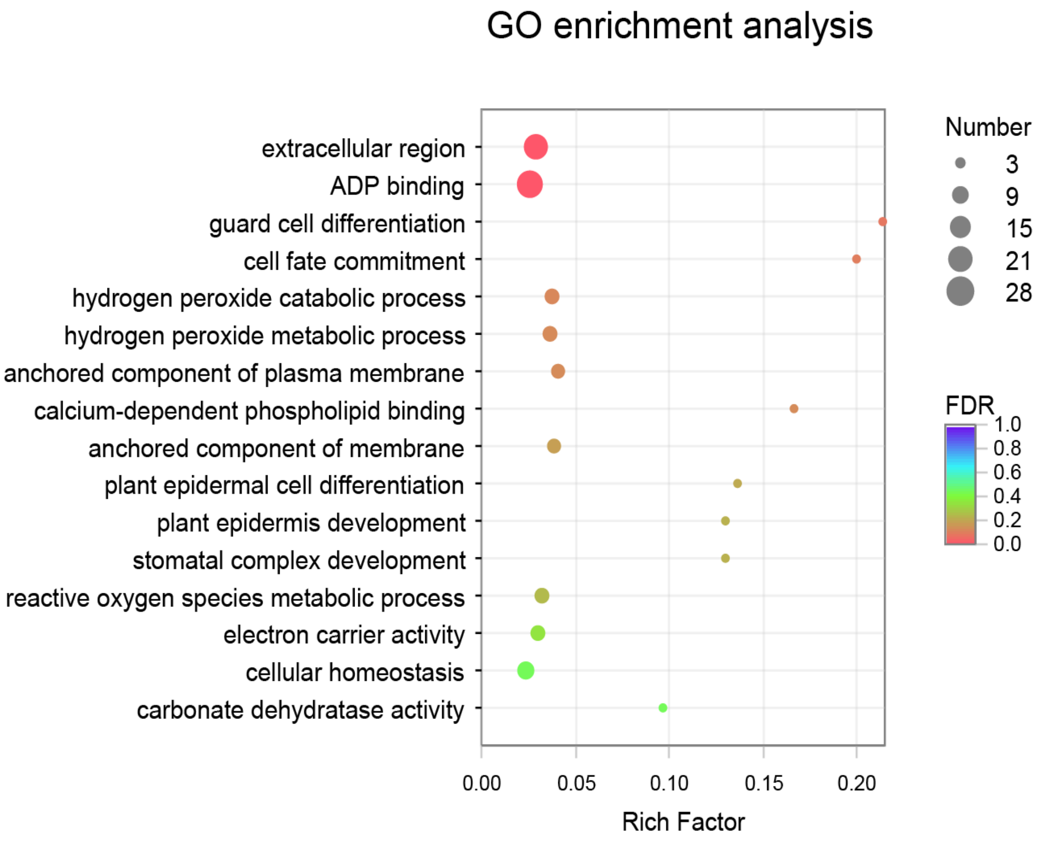


**Figure S6.** Go enrichment of the expanded gene families of *P. villosa*

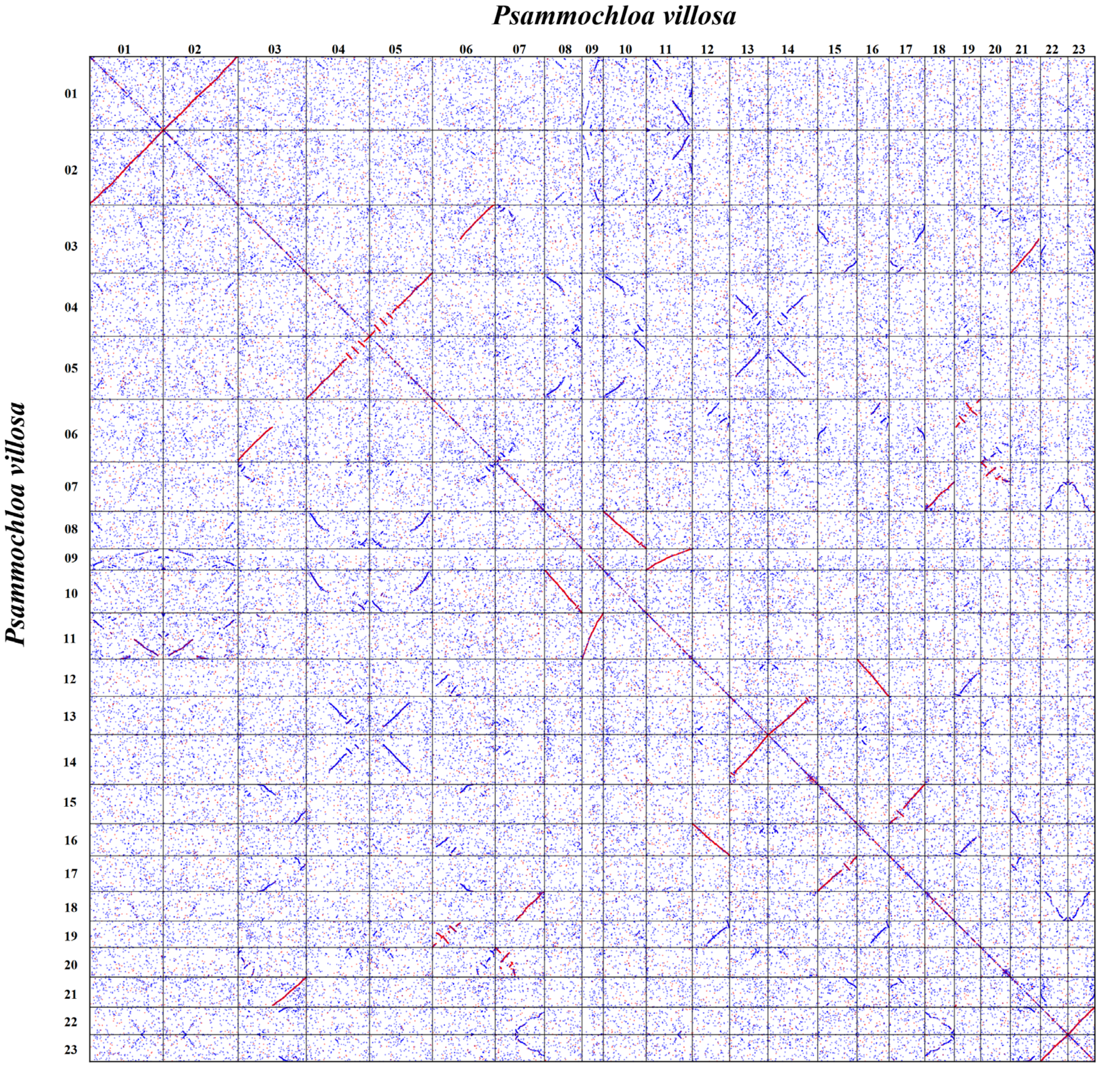


**Figure S7.** Overview of dotplots within the *P. villosa* genome for paralogous genes. Best-hit (orthologous) genes are red dots, secondary hits are blue dots, and the others are shown in gray. Highlights show the best matched chromosomal regions.

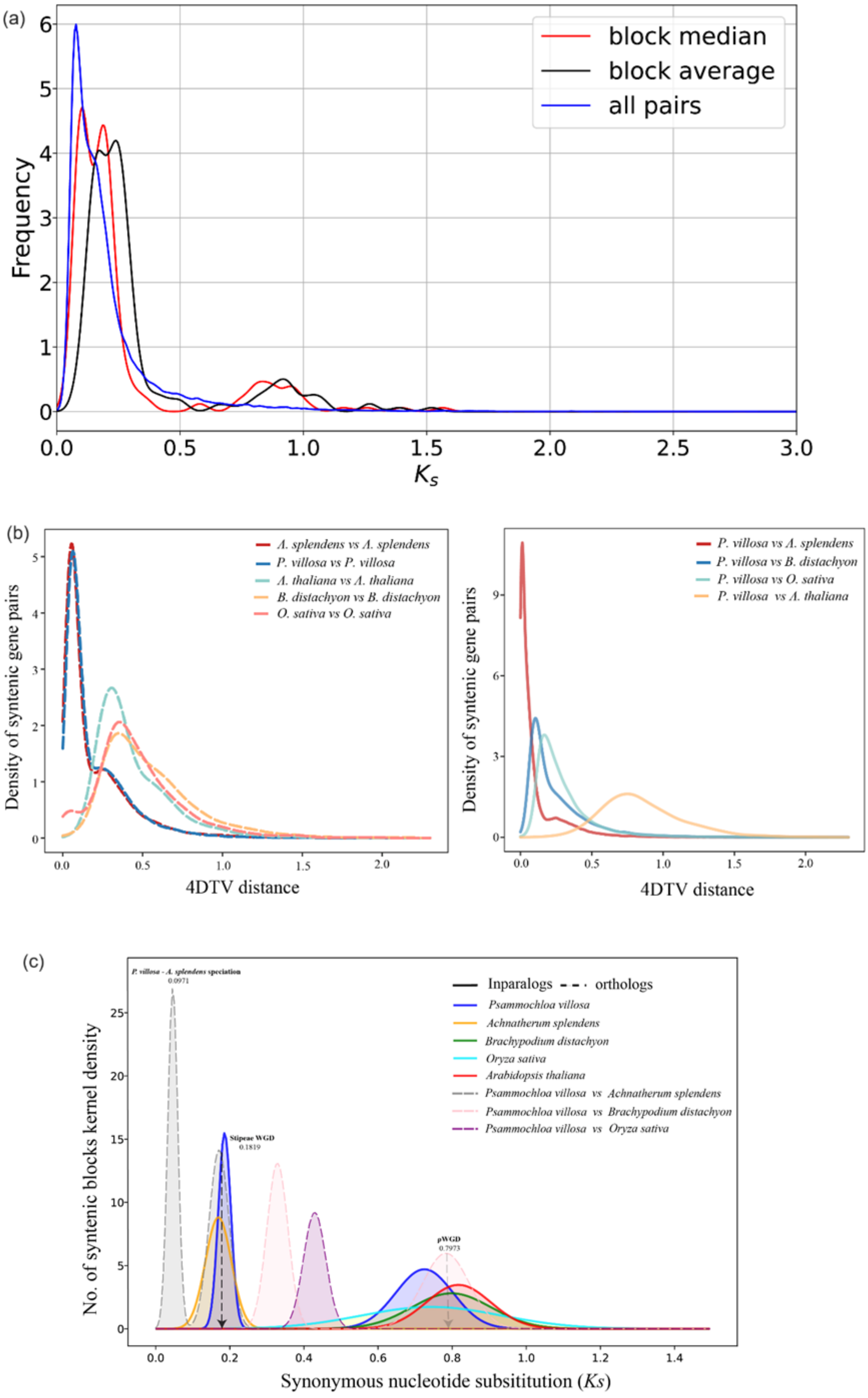


**Figure S8.** Collinearity analysis of interspecies genome based on frequency of synonymous substitution levels (*Ks* values and 4DTV) of syntenic orthologous. (a) *Ks* peak of *P. villosa* with gaussian distribution. (b) *Ks* pair in syntenic block among different species, orthologs (dashed curves) and paralogous genes (solid curves) respectively corresponded to specie's genome duplication and differentiation after evolutionary rate correction. (c) Estimation of recent WGD and divergence time based on distribution of synonymous substitution (*Ks*) and hypothesis of neutral rate of evolution (T = *Ks*/*2r*).

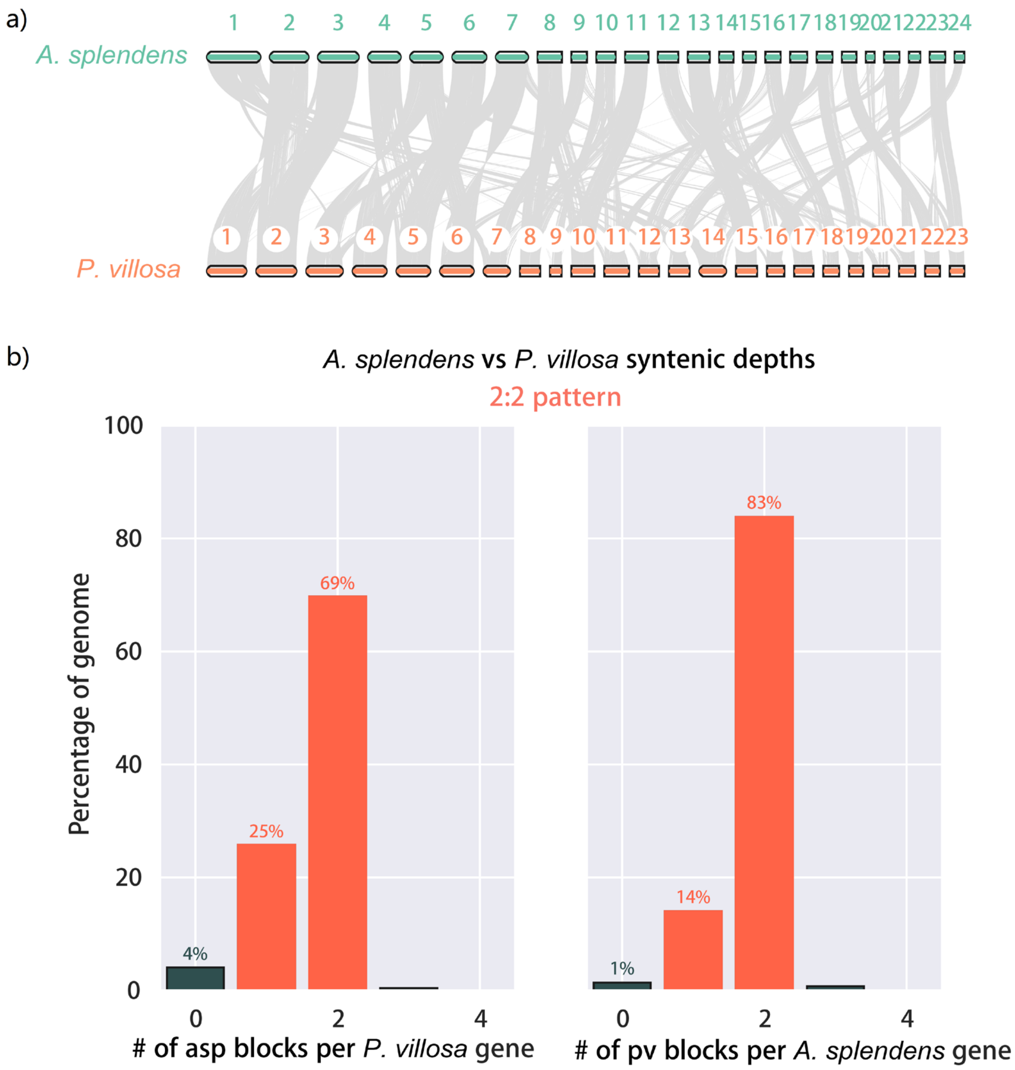


**Figure S9.** A typical collinearity pattern between genomic regions of *P. villosa* and *A. splendens*. (a) Synteny block where grey lines represent one-to-one gene synteny (b) depth of collinearity of two genes of *P. villosa* corresponding to two genes of *A. splendens*.

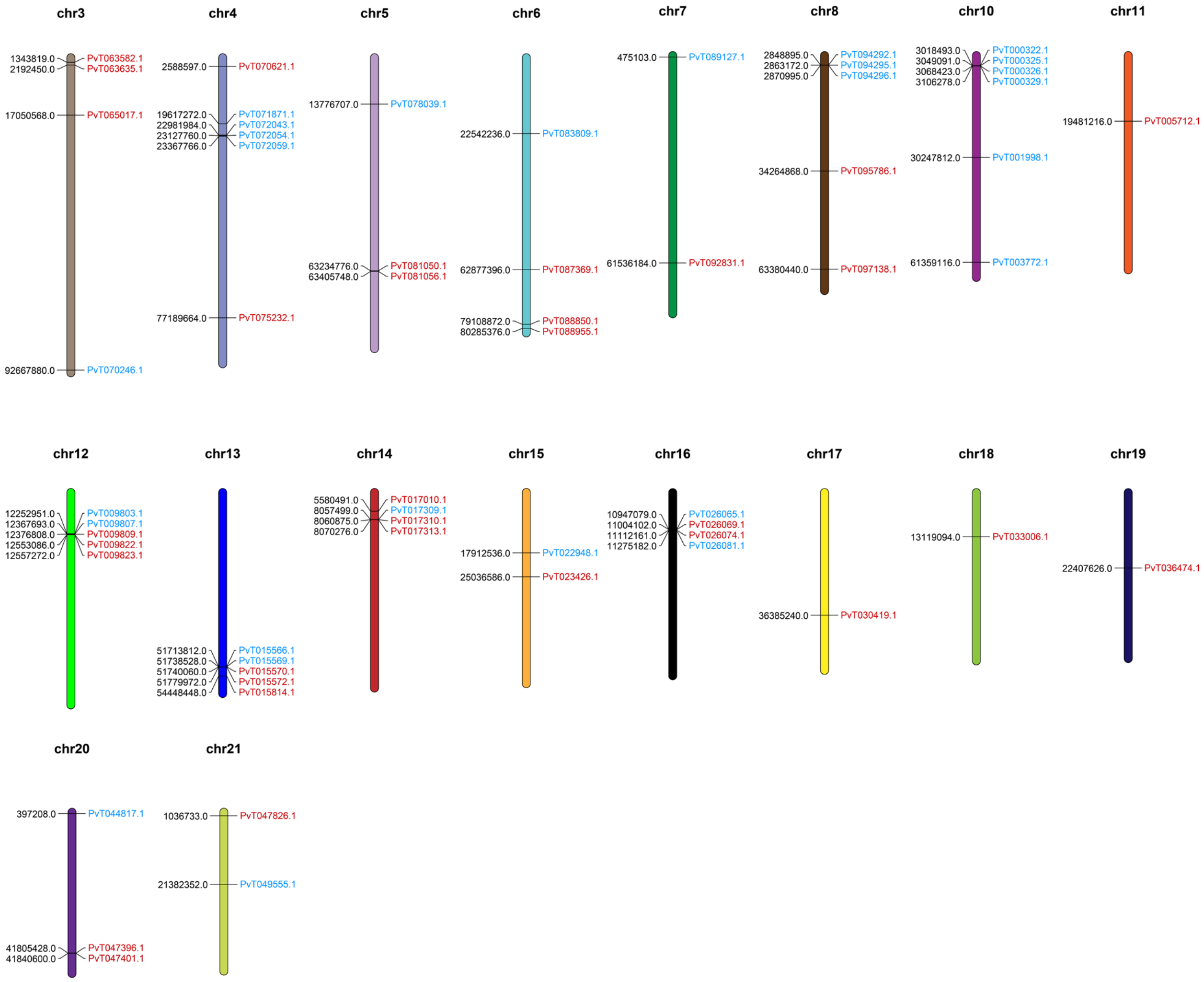


**Figure S10.** Distribution of all *XTHs* genes on chromosomes of *P. villosa*.

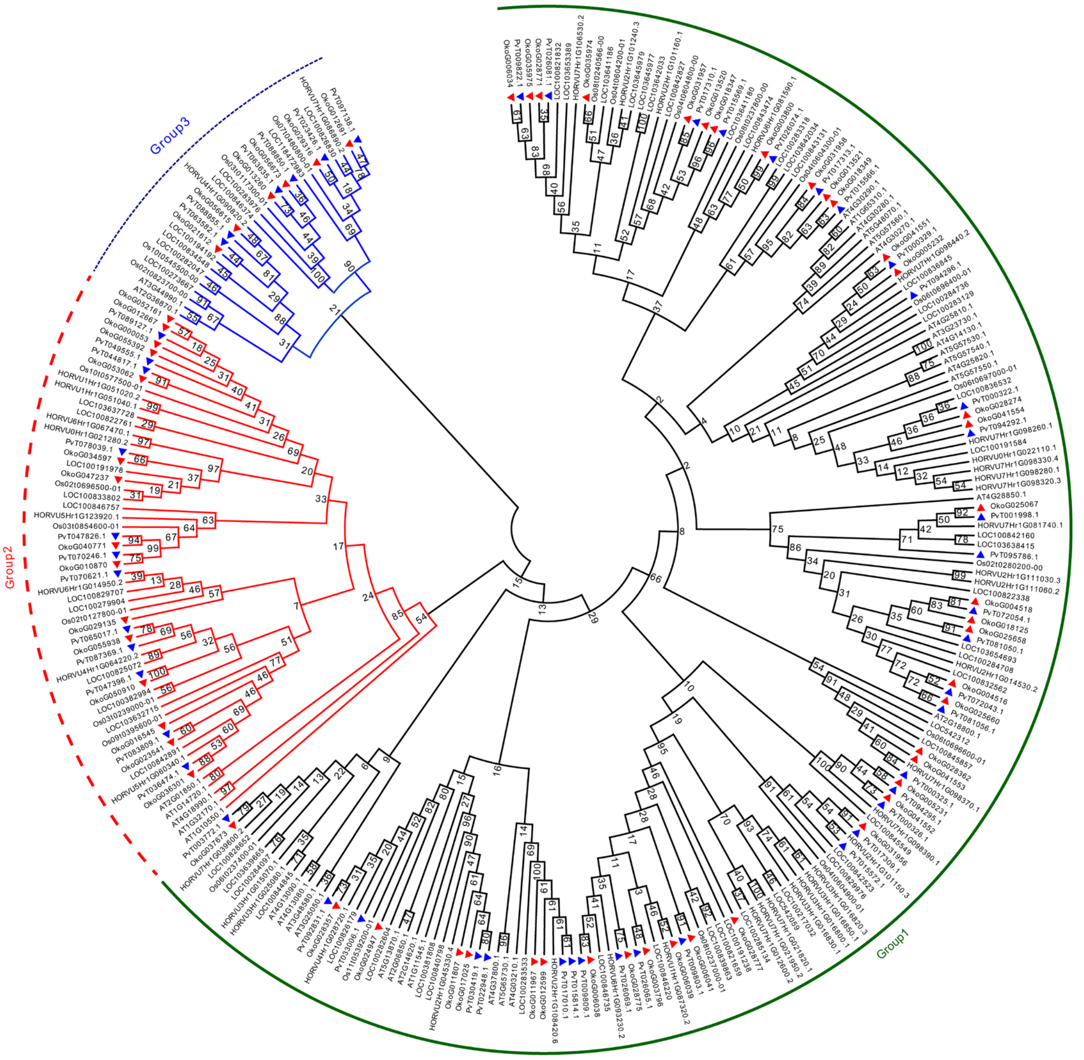


**Figure S11.** The phylogenetic relationship of *XTHs* gene derived from *P. villosa*, *O. sativa*, *B. distachyon*, *H. vulgare*, and *N. splendens*. Red line indicates class III orthologues experience expansion during *P. villosa* evolution.

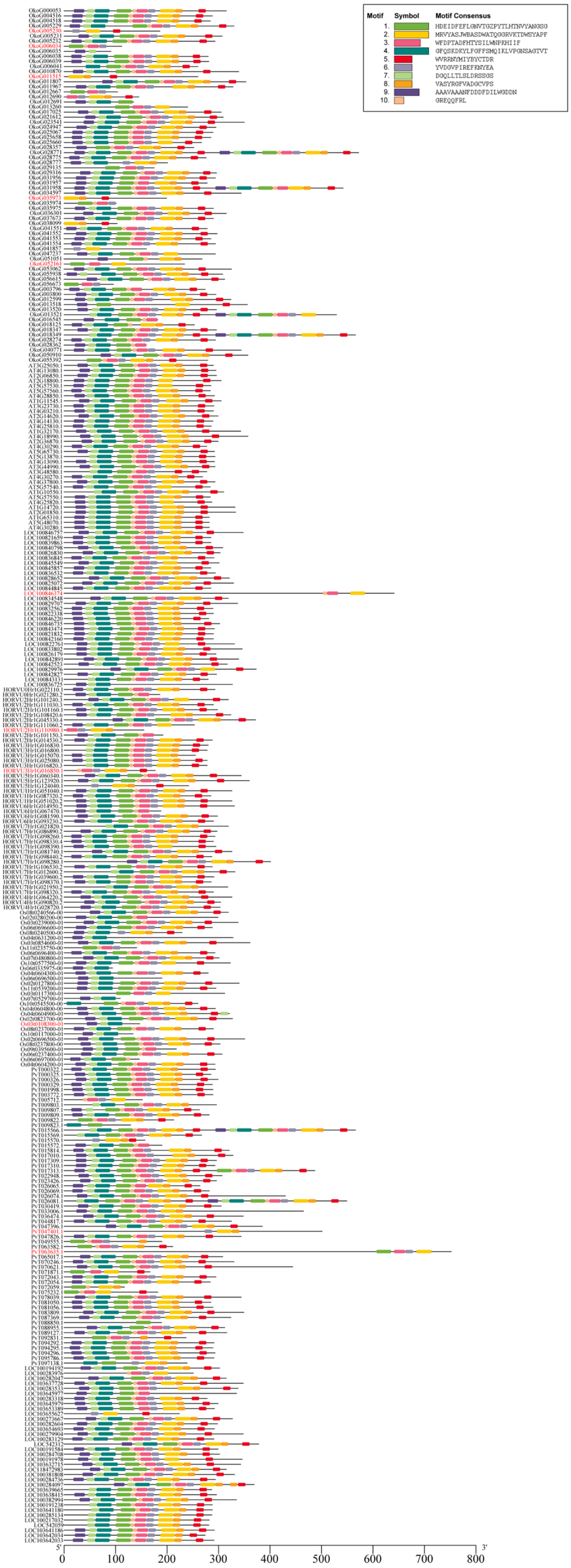


**Figure S12.** Structural analysis of the ten conserved motifs in *XTHs* gene structures.

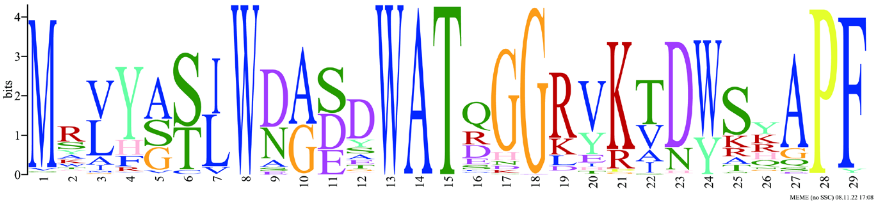

**Figure S13.** Sequence composition of conserved motif 2 of *P. villosa*.

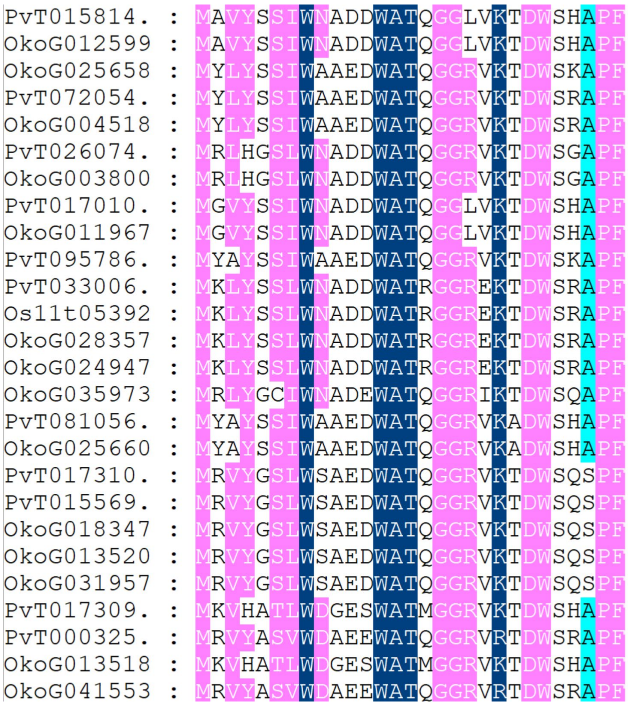


**Figure S14.** Multiple alignment of motif 2 of *XTHs* for showing the conserved secondary structures. The conserved residues were shown in blue frames, among which the identity residues were indicated by white letters in red boxes, and the similar residues by black letters in pink boxes.

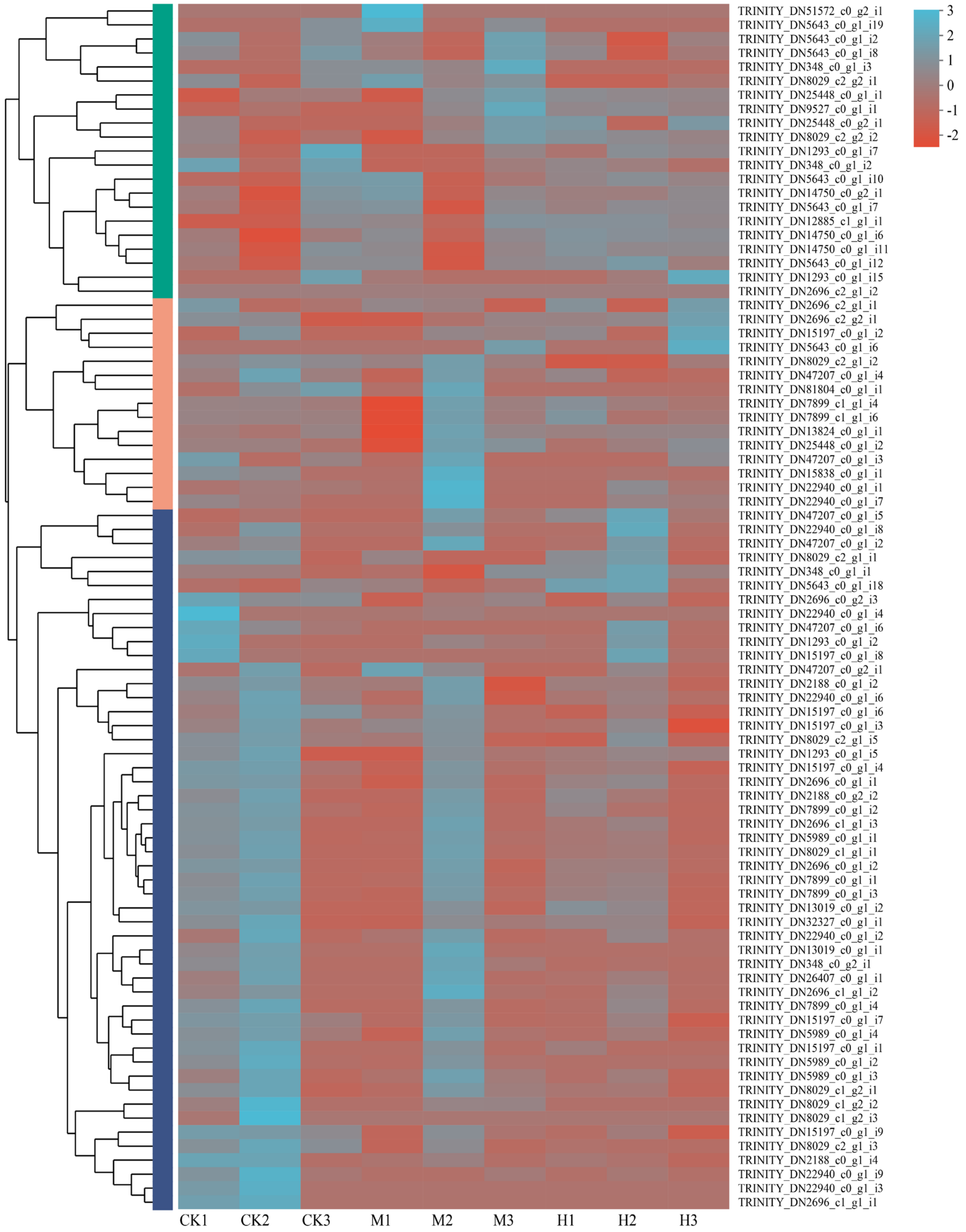


**Figure S15.** Gene expression of the xyloglucan endo-transglucosylase/hydrolase family (59 *XTHs*) under different drought treatments using 10% (TM) and 20% (TH) PEG-6000.

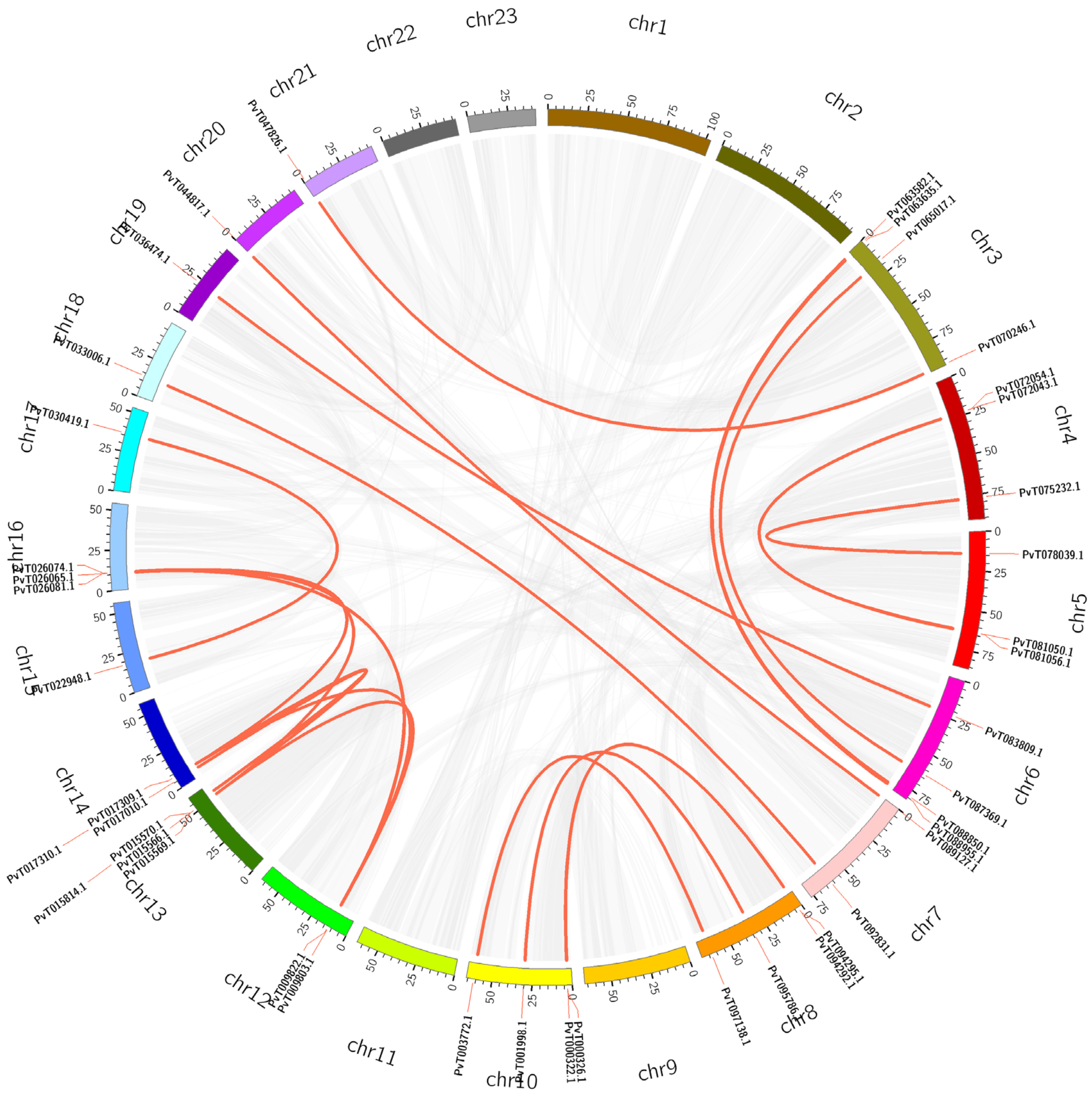

**Figure S16.** Genomics collinearity of the *PvXTHs* family (59 genes) by synteny of *P. villosa*.

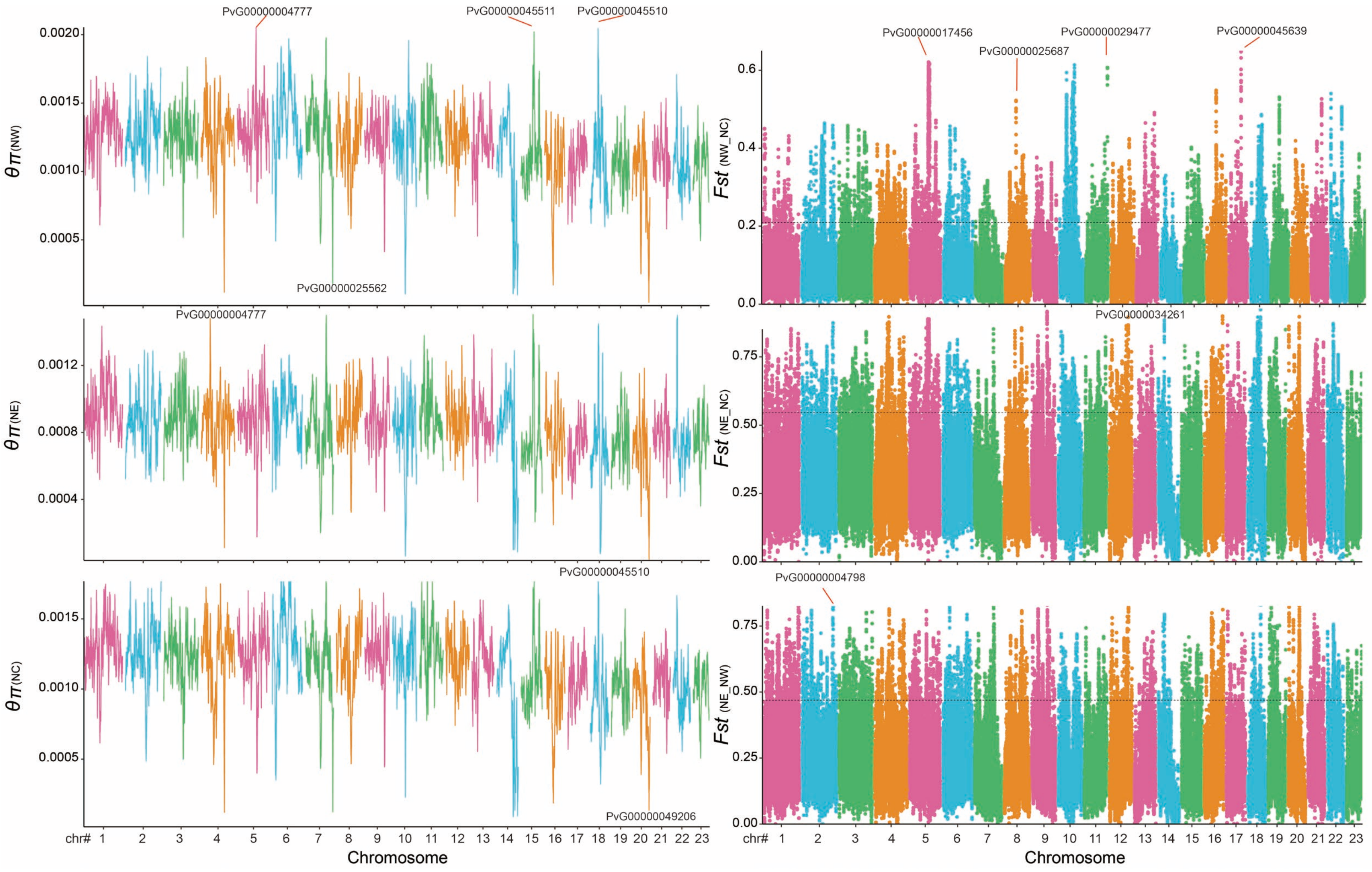


**Figure S17.** Genetic diversity, population divergence between northwest and northeast lineages.

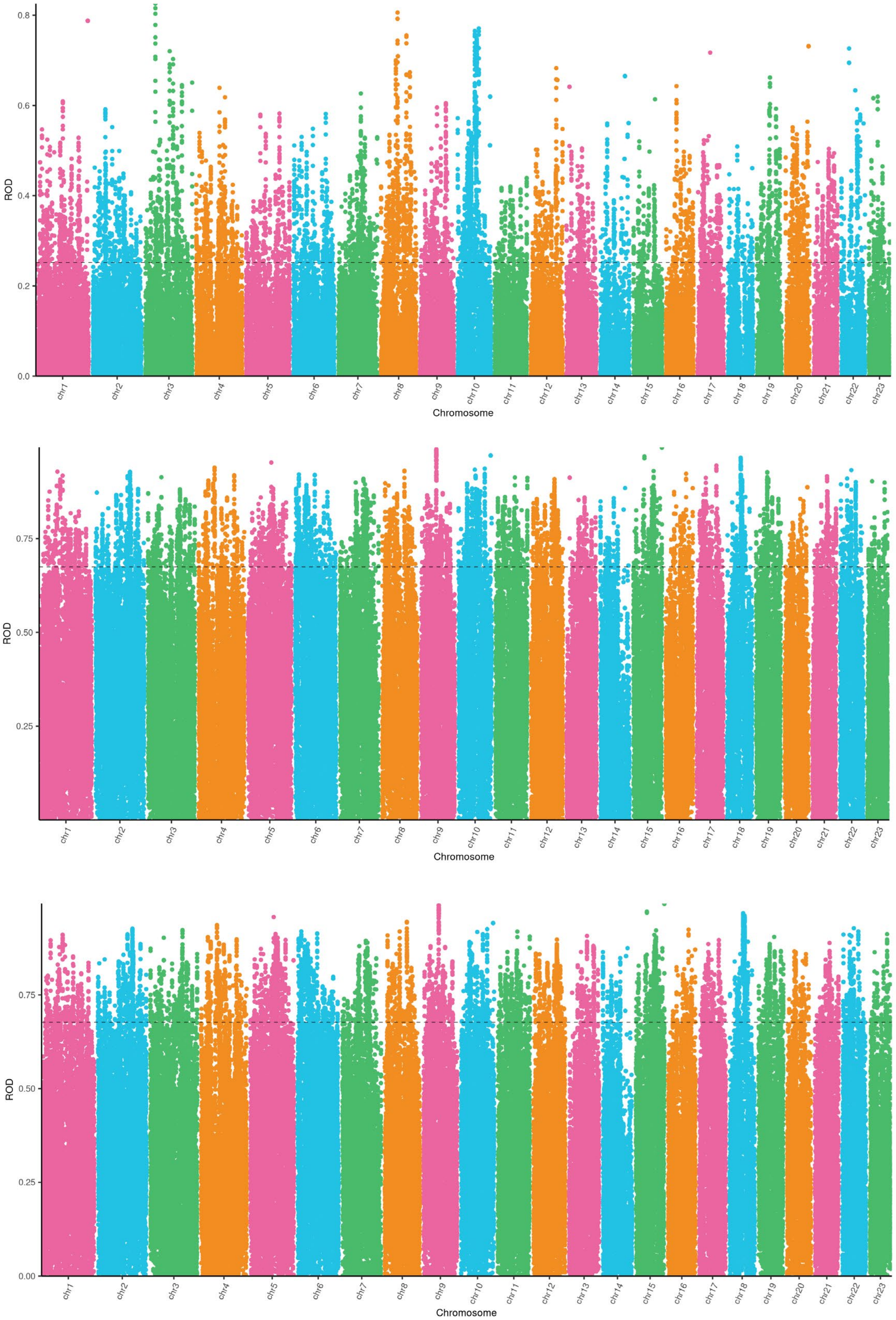


**Figure S18**. Genome-wide scanning to identify selection region by the reduction of diversity (ROD) northwest and northeast lineages.

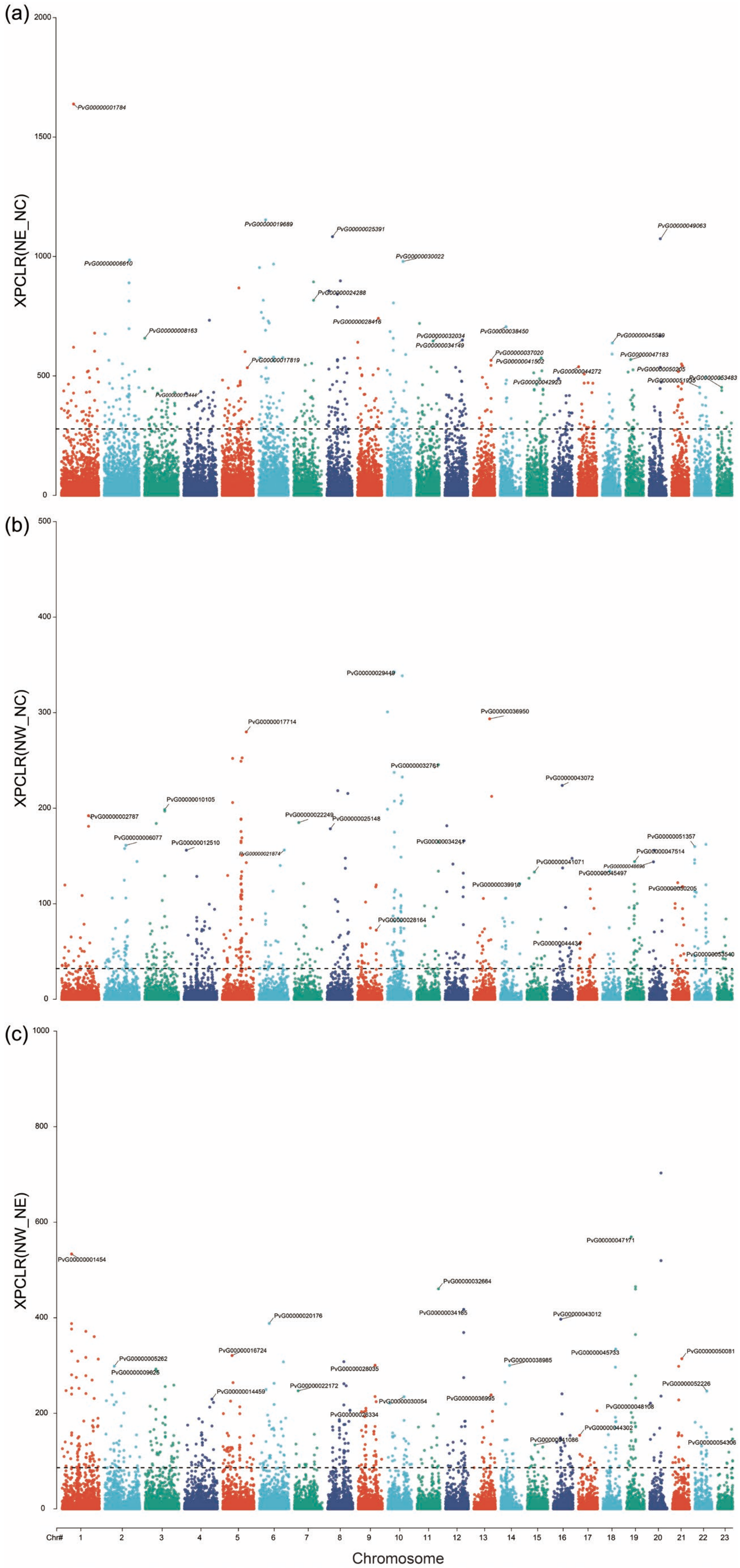

**Figure S19.** The estimation of XP-CLR score among northwest, northeast and northcentral lineages.

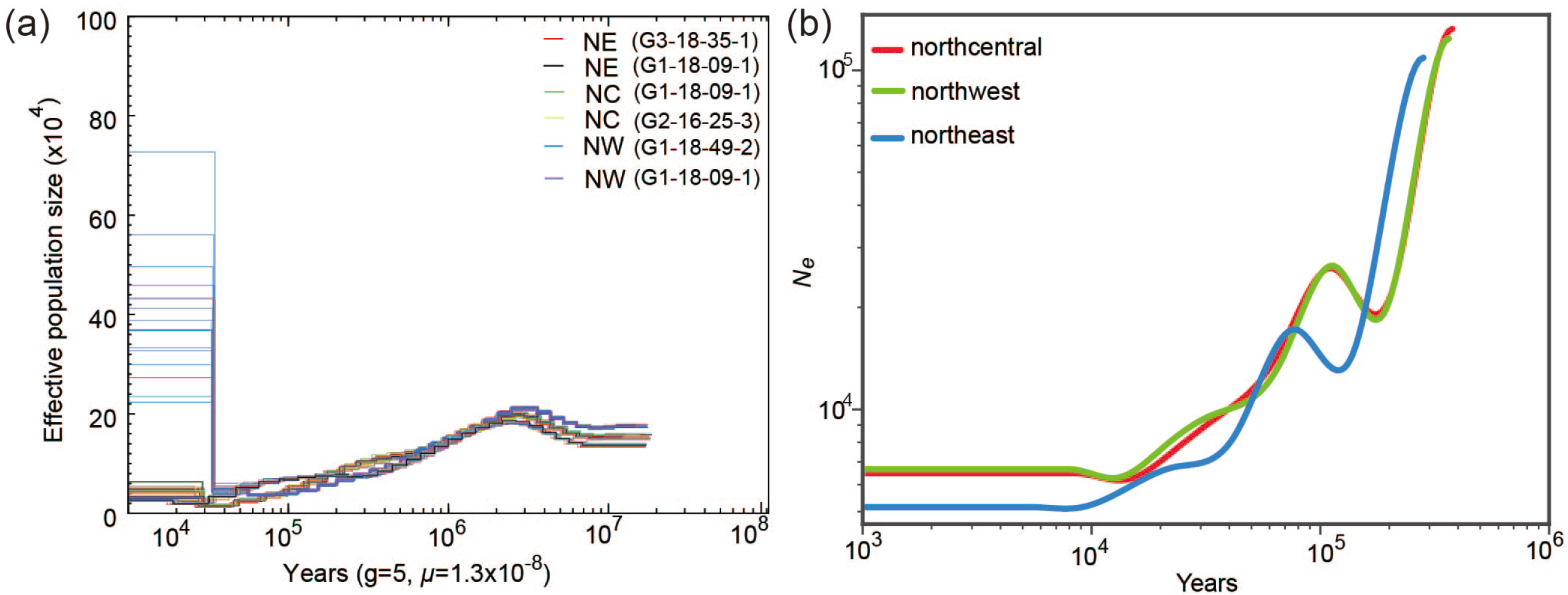


**Figure S20**. Demographic history and effective population size of *P. villosa*. (a) Changes in effective population size (*Ne*) through time inferred by the Pairwise Sequentially Markovian Coalescent (PSMC) model. (b) Changes in effective population size (*Ne*) through time inferred by SMC++ model.

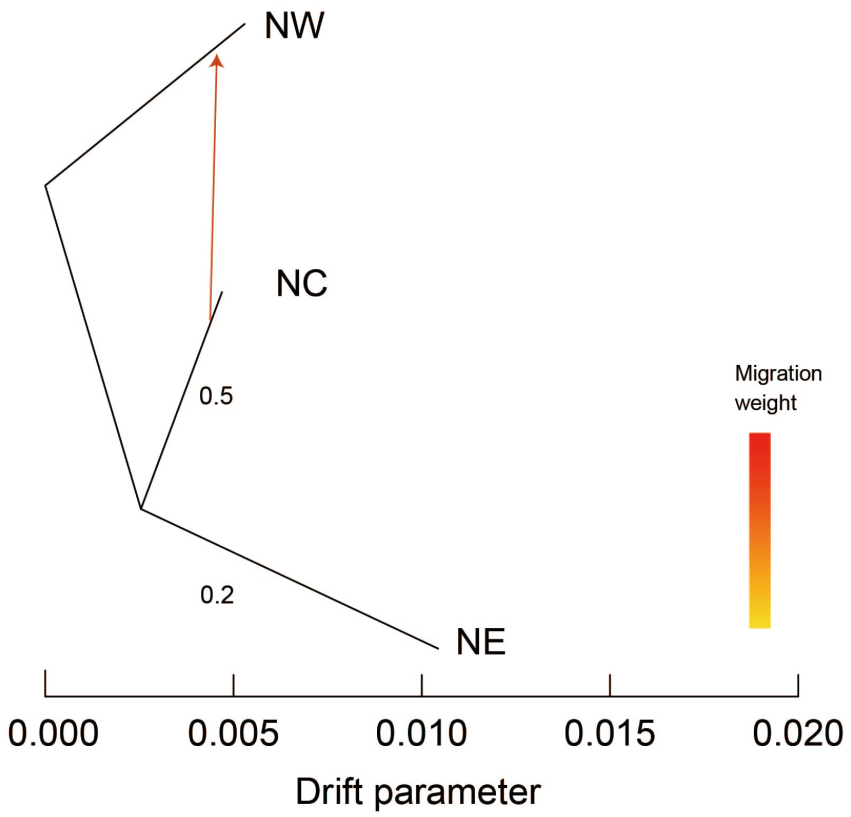


**Figure S21**. Trees inferred by TreeMix. Gene flow test generated 126 independent simulations of 23 chromosomes from each population using the demographic model without migration, and inferred population trees.

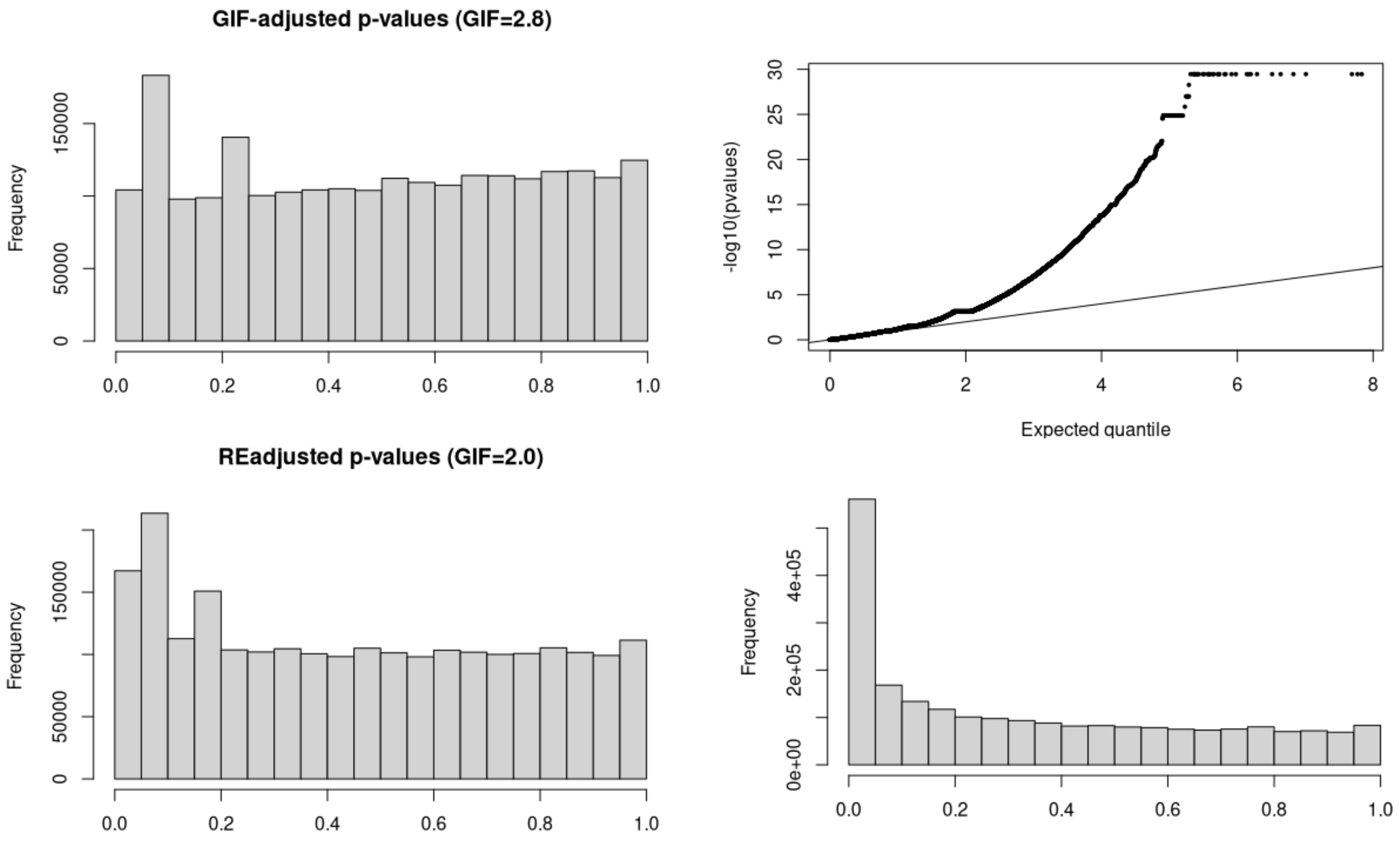


**Figure S22.** Mapping the location of the 324,223 environmental adaptive genes detected by LFMM model across the whole genome.

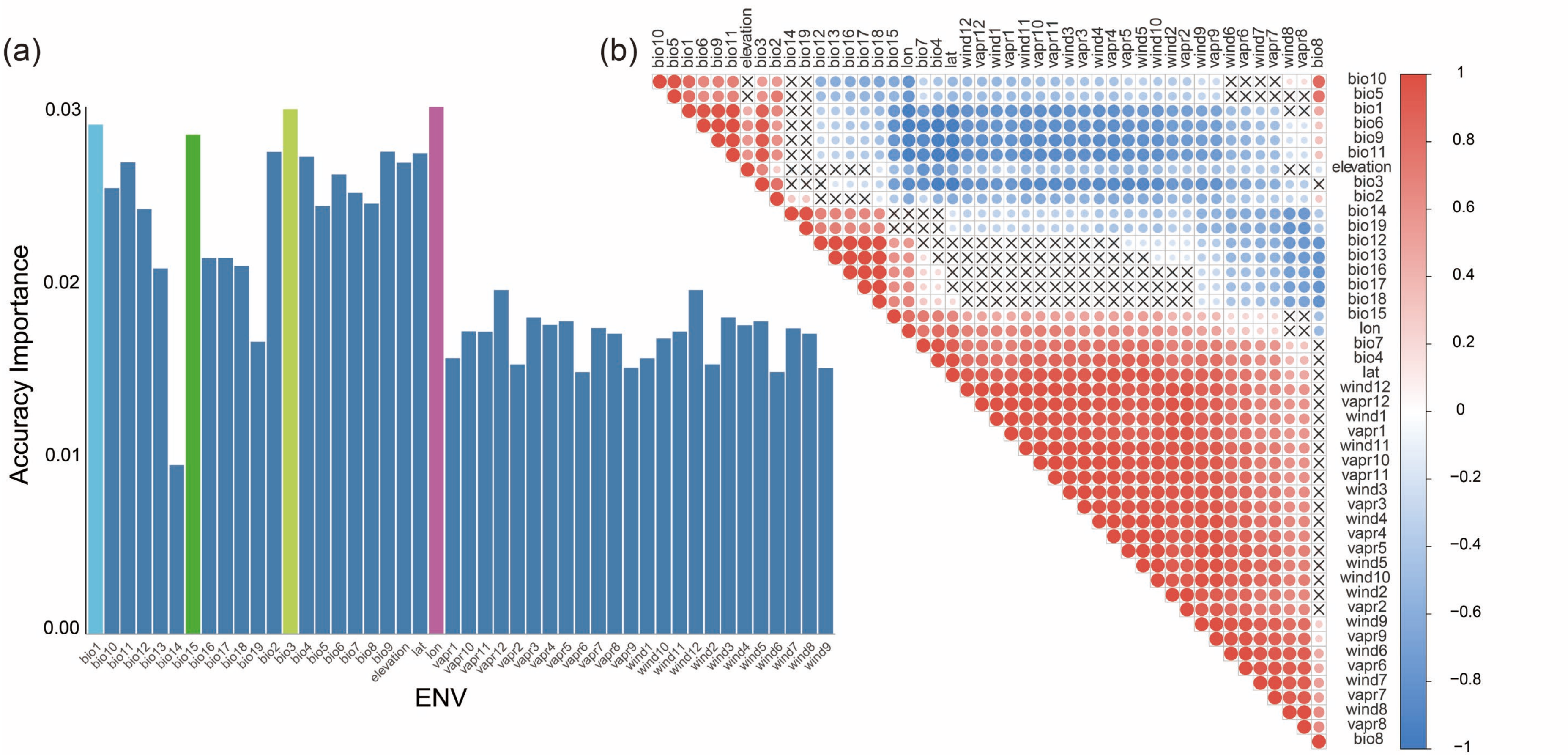


**Figure S23**. The sort of accuracy importance for each variate's contribution. (a) based on the gradient forest (GF) analysis ranking to model the turnover in allele frequencies along present environmental gradients and (b) correlation of 46 climatic and environmental variates.

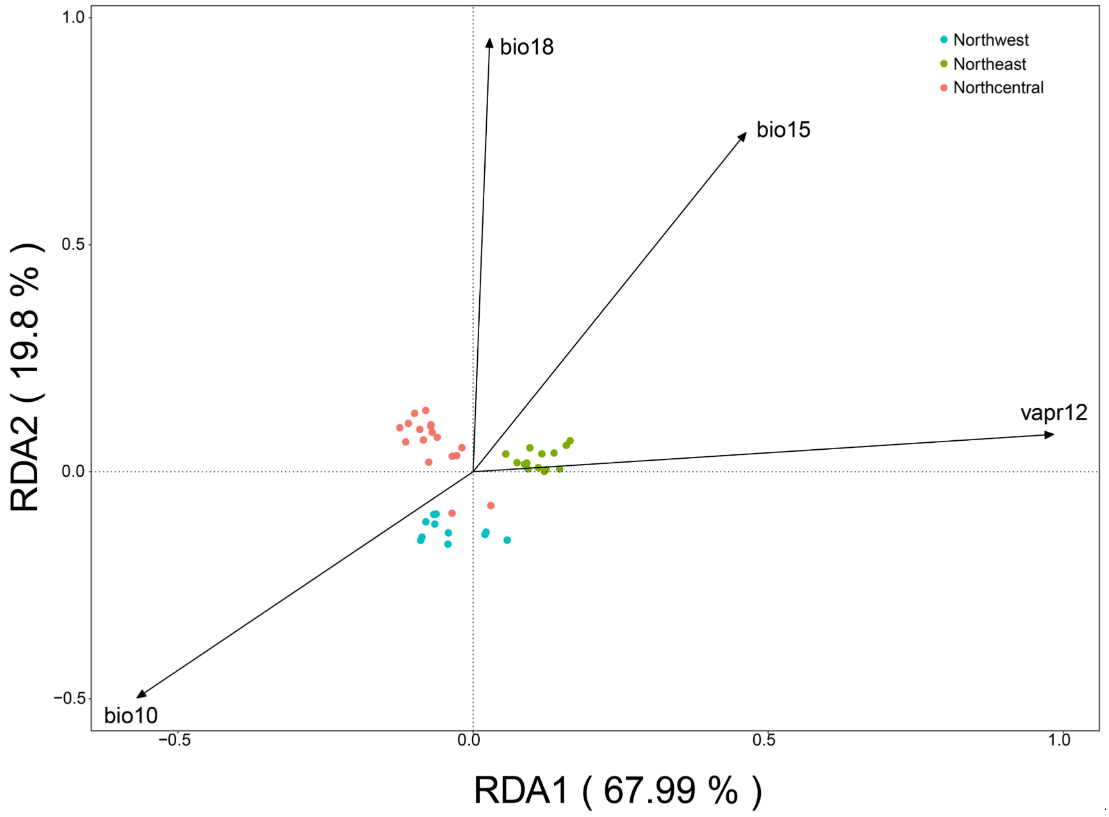


Figure S24. The redundancy analysis (RDA) selected the four variables associated with population genetic variation.
